# SpeciesRax: A tool for maximum likelihood species tree inference from gene family trees under duplication, transfer, and loss

**DOI:** 10.1101/2021.03.29.437460

**Authors:** Benoit Morel, Paul Schade, Sarah Lutteropp, Tom A. Williams, Gergely J. Szöllősi, Alexandros Stamatakis

## Abstract

Species tree inference from gene family trees is becoming increasingly popular because it can account for discordance between the species tree and the corresponding gene family trees. In particular, methods that can account for multiple-copy gene families exhibit potential to leverage paralogy as informative signal. At present, there does not exist any widely adopted inference method for this purpose. Here, we present SpeciesRax, the first maximum likelihood method that can infer a rooted species tree from a set of gene family trees and can account for gene duplication, loss, and transfer events. By explicitly modelling events by which gene trees can depart from the species tree, SpeciesRax leverages the phylogenetic rooting signal in gene trees. SpeciesRax infers species tree branch lengths in units of expected substitutions per site and branch support values via paralogy-aware quartets extracted from the gene family trees. Using both empirical and simulated datasets we show that SpeciesRax is at least as accurate as the best competing methods while being one order of magnitude faster on large datasets at the same time. We used SpeciesRax to infer a biologically plausible rooted phylogeny of the vertebrates comprising 188 species from 31612 gene families in one hour using 40 cores. SpeciesRax is available under GNU GPL at https://github.com/BenoitMorel/GeneRax and on BioConda.

## Introduction

Phylogenetic species tree inference constitutes a challenging computational problem. Accurate and efficient tools for species tree inference exhibit a substantial potential for obtaining novel biological insights.

The concatenation or supermatrix approach has long been the gold standard for species tree inference. Here, gene sequences are first aligned and subsequently concatenated into a single, large supermatrix. Then, statistical tree inference methods (maximum likelihood (Kozlov *et al*., 2019; Minh *et al*., 2020) or Bayesian inference (Aberer *et al*., 2014; Ronquist *et al*., 2012)) are applied to infer a tree on these supermatrices. The concatenation approach heavily relies on accurate orthology inference, which still constitutes a challenging problem (Altenhoff *et al*., 2019). In addition, concatenation methods were shown to be statically inconsistent under the multispecies coalescent model (Kubatko and Degnan, 2007; Mendes and Hahn, 2017) because of potential incomplete lineage sorting (ILS).

As gene family tree (GFT) methods can alleviate some of the pitfalls of the supermatrix approach, they are becoming increasingly popular. GFT methods can take into account that the evolutionary histories of the gene trees and the species tree are discordant due to biological phenomena such as ILS, gene duplication, gene loss, and horizontal gene transfer (HGT).

At present, the most commonly used GFT tools (Bouckaert *et al*., 2014; Zhang *et al*., 2017) only model ILS and are limited to single-copy gene families. These methods also heavily rely on accurate orthology inference and discard large amounts of potentially informative data. Approaches that can handle multiple-copy gene families exist, but have not been widely adopted yet (Boussau *et al*., 2012; Molloy and Warnow, 2020; Wehe *et al*., 2008; Zhang *et al*., 2019). Here, we focus on describing, evaluating, and making available a novel method for inferring reliable species trees from multiple-copy gene families in the presence of both paralogy and HGT. For instance, HGT is particularly challenging when analysing microbial clades, because supermatrix analyses can be misled in unpredictable ways if HGTs are included in the concatenation (Dombrowski *et al*., 2020; Williams and Embley, 2014).

One class of existing methods to infer species trees from multiple-copy gene families attempts to simultaneously estimate the GFTs and the species tree (Boussau *et al*., 2012; de Oliveira Martins and Posada, 2017). However, these methods are computationally demanding and are limited to small datasets comprising less than 100 species.

Another class of existing methods handles the GFT inference and the species tree inference steps separately. As input they require a set of given, fixed GFTs and do not attempt to correct the GFTs during the species tree inference step. DupTree (Wehe *et al*., 2008) and DynaDUP (Bayzid *et al*., 2013) search for the species tree with the least parsimonious reconciliation cost, measured as the number of duplication events in DupTree, and the sum of duplication and loss events in DynaDUP. STAG (Emms and Kelly, 2018) infers a species tree by applying a distance method to each gene family that covers *all* species, and subsequently builds a consensus tree from all these distance-based trees. However, STAG ignores a substantial fraction of signal by discarding gene families that do not cover all species. FastMulRFS (Molloy and Warnow, 2020) extends the definition of the Robinson-Foulds (RF) distance to multiple-copy GFTs and strives to minimize this distance between the species tree and all input GFTs. More recently, with ASTRAL-Pro (Zhang *et al*., 2019) a promising improvement of ASTRAL was released to handle multiple-copy GFTs: ASTRAL-Pro uses dynamic programming to infer the species tree that maximizes a novel measure of quartet similarity that accounts for orthology and paralogy. All of the above methods are non-parametric and do not deploy a probabilistic model of evolution. In addition, none of them explicitly models HGT.

Here, we present SpeciesRax, the first maximum likelihood method for inferring a rooted species tree from a set of GFTs in the presence of gene duplication, gene loss, and HGT. We implemented it in the GeneRax framework, our recently published species-tree-aware GFT correction tool (Morel *et al*., 2019). SpeciesRax takes as input a set of multiple sequence alignments (MSAs) and/or a set of GFTs. If MSAs are provided, SpeciesRax will infer one maximum likelihood GFTs tree per gene family using RAxML-NG (Kozlov *et al*., 2019). Thereafter, SpeciesRax first generates an initial, reasonable (i.e., non-random) species tree by applying MiniNJ (which we also introduce in this paper), our novel *distance* based method for species tree inference from GFTs in the presence of paralogy. MiniNJ shows similar accuracy as other non-parametric methods while being at least two orders of magnitude faster on large datasets. Finally, SpeciesRax executes a maximum likelihood tree search heuristic under an explicit statistical gene loss, gene duplication, and HGT model starting from the MiniNJ species tree. When the species tree search terminates, SpeciesRax calculates approximate branch lengths in units of mean expected substitutions per site. Furthermore, it quantifies the reconstruction uncertainty by computing novel quartet-based branch support scores on the species tree. Since we implemented all of these new methods in our GeneRax software, users can now perform GFT inference, species tree inference, GFT correction, and GFT reconciliation with the species tree using a single tool. We show that SpeciesRax is fast and at least as accurate as the best competing species tree inference tools. In particular, SpeciesRax is twice more accurate (in terms of relative RF distance to the true species trees) than all other tested methods on simulations with large numbers of paralogous genes.

## Method

We recently (Morel *et al*., 2019) introduced the *undatedDTL* model that describes the evolution of a GFT along a species tree through gene duplication, gene loss, speciation, and HGT events. In addition, we described an algorithm for computing the corresponding reconciliation likelihood, that is, the probability of observing a set of GFTs *𝒢* =(*G*_1_,…,*G*_*n*_) given a rooted species tree *S* and the set *N* of duplication, loss, and HGT intensities:

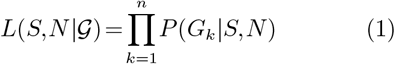

As already mentioned, SpeciesRax takes a set of unrooted GFTs as input. It starts its computations from an initial species tree that can either be randomly generated, user-specified, or inferred using our new distance method MiniNJ. Then, it performs a tree search for the rooted species tree *S* and the model parameters *N* that maximize the reconciliation likelihood *L*(*S,N* |*𝒢*). At the end of the search, it also calculates support values for the inner branches of the inferred species tree from the GFTs. Finally, we also describe the adaptation of our likelihood score to better account for missing data and inaccurate assignment of sequences to gene family clusters. Computing a reasonable initial species tree with MiniNJ

Here, we introduce MiniNJ (Minimum internode distance Neighbor Joining), our novel distance based method for inferring an unrooted species tree in the presence of paralogy. MiniNJ is fast, that is, it is well-suited for generating an initial species tree for the subsequent maximum likelihood optimization. MiniNJ is inspired by NJst (Liu and Yu, 2011), a distance based method that performs well in the *absence* of paralogy. Initially, we briefly outline the NJst algorithm, and subsequently describe our modifications.

NJst initially computes a distance matrix from the unrooted GFTs and then applies Neighbor Joining (NJ) to reconstruct the species tree. NJstdefines the gene internode distance *D*_*g*_ such that *D*_*g*_(*x,y*) is the number of internal nodes between the terminal nodes *x* and *y* in a GFT. NJst computes the distance between two species as the average over the internode distances between all pairs of gene copies mapped to those two species.

More formally, let *a* and *b* be two species. Let *K* be the number of GFTs. Let *m*_*ak*_ be the terminal nodes from the GFT *k* mapped to species *a*. Let *x*_*iak*_ be the *i*th terminal node from the GFT *k* mapped to species *a*. NJst defines the distance matrix *D*_*NJst*_ as follows:

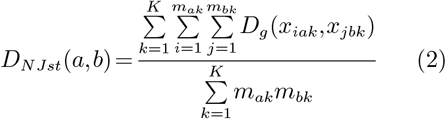

NJst has two drawbacks. First, it accounts for all pairs of gene copies, including paralogous gene copies that do not contain information about speciation events (see Fig. 1). Secondly, it assigns very high (quadratic) weights to gene families comprising a high number of gene copies: for instance, a gene family *k*_1_ with 5 gene copies in both species *a* and *b* will contribute 25 times to the distance between *a* and *b*, while a single-copy family *k*_2_ will only contribute once. For instance, 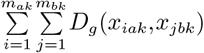 is the sum over 25 gene internode distances for family *k*_1_ and of only one gene internode distance for family *k*_2_.

**FIG. 1.**
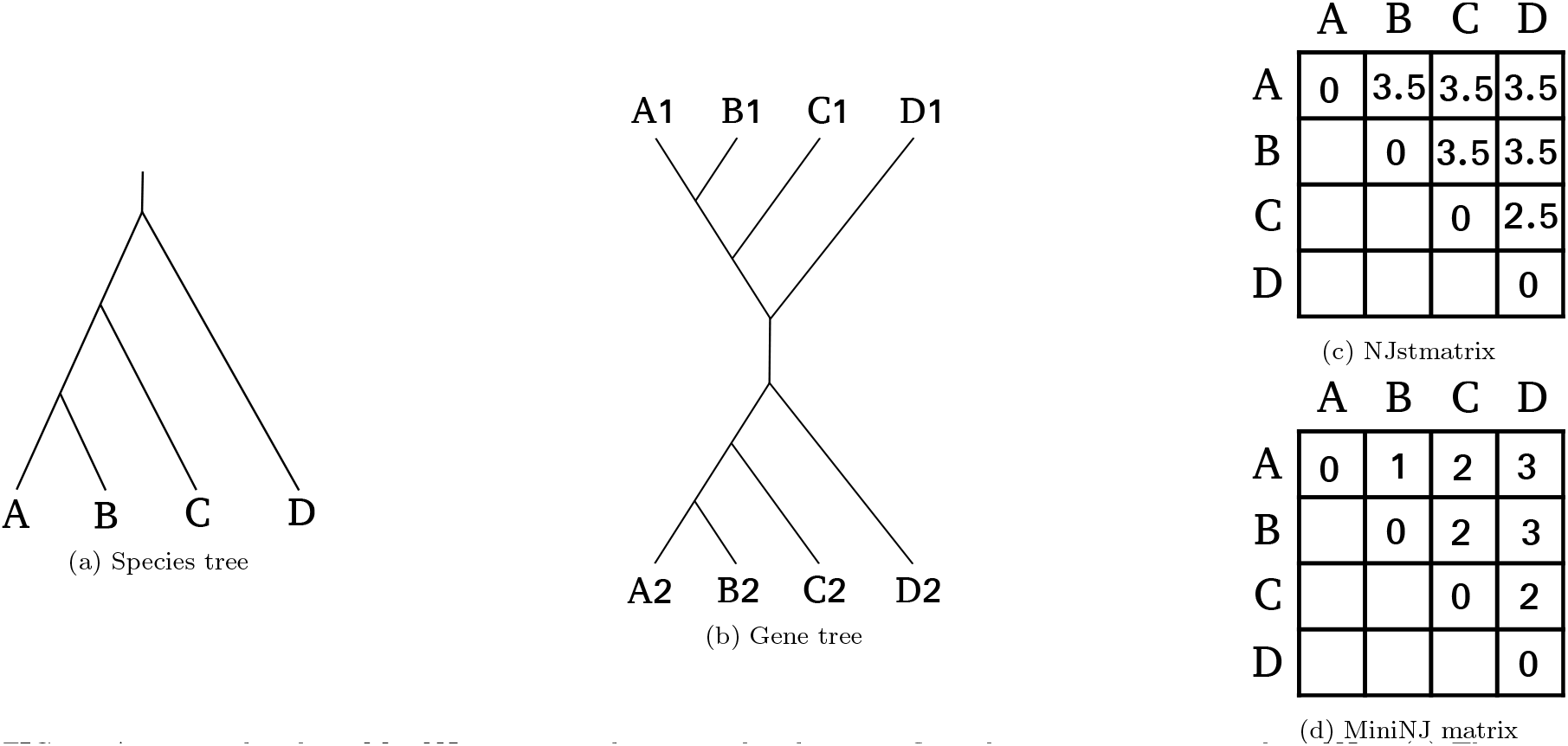
An example where MiniNJ computes distances that better reflect the true species tree than NJst. (a) The true rooted species tree. (b) A GFT resulting from a duplication at the root of the species tree. (c) The distance matrix *D*_*NJst*_ computed with NJst, incorrectly suggesting that all species are equidistant, apart from *C* and *D*. This is the result of distance overestimation due to paralogous genes: for instance, species *A* and *B* are neighbors in the species tree, but the genes *A*2 and *B*1 are very distant from each other in the gene tree, because they start diverging from an early duplication event (paralogous genes). (d) Distance matrix *D*_*MiniNJ*_ computed with MiniNJ. The gene internode distances correctly reflect the species distances, because MiniNJ successfully pruned pairs of paralogous genes, such as *A*2 and *B*1, and only accounted for orthologous genes, such as *A*1 and *B*1.

**FIG. 2.**
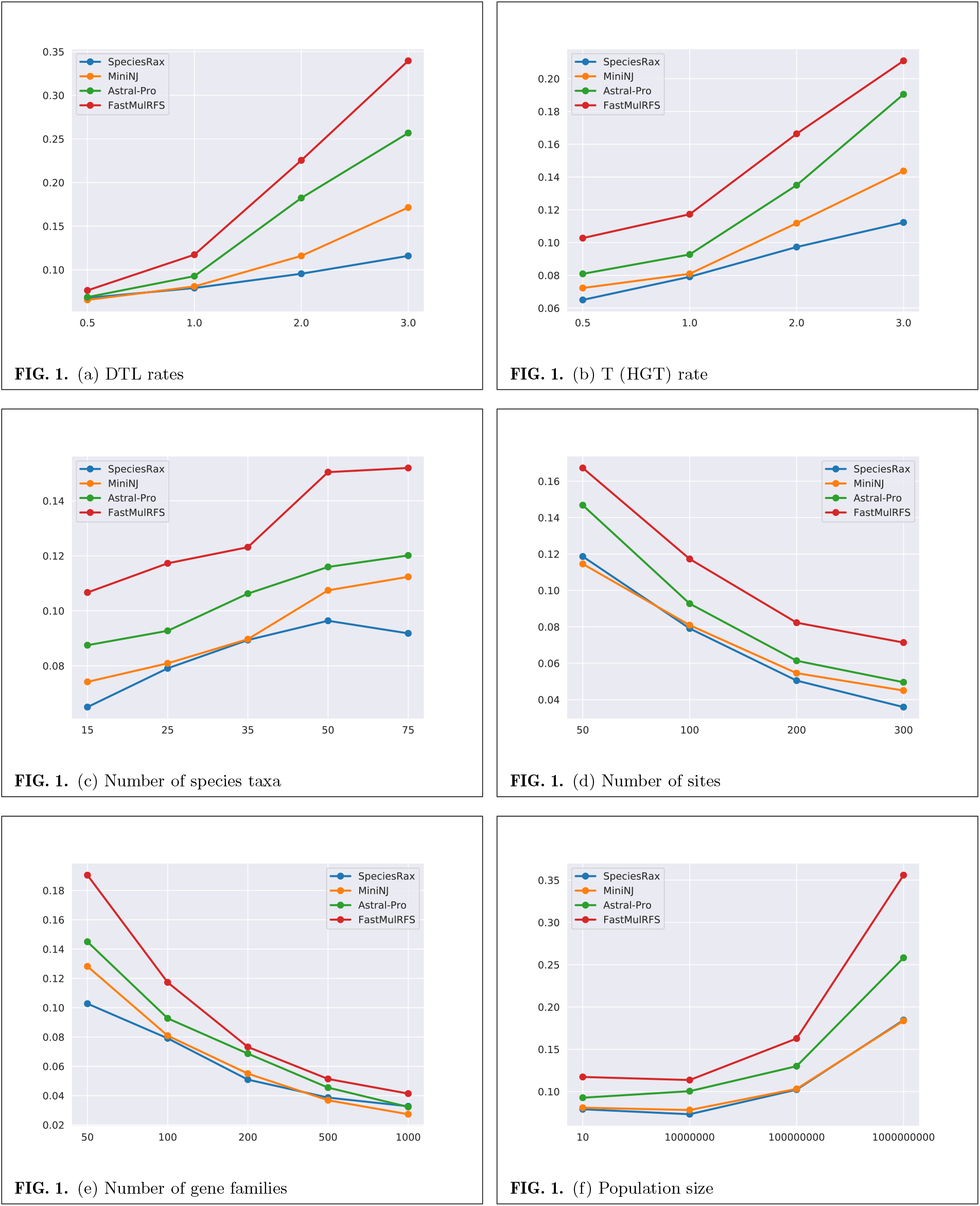
Average unrooted RF distance between inferred and true species trees, in the presence of duplication, loss and transfers.

Since the normalization by the number of gene internode distances is conducted after summing over all these quantities (with the denominator in Eq 2), the contributions of families *k*_1_ and *k*_2_ are unbalanced.

MiniNJ adapts Eq. 2 to address these two issues. It attempts to discard pairs of paralogous gene copies by only considering the two closest GFT terminal nodes mapped to a pair of species for each family, according to the internode distance: let *δ*_*abk*_ be equal to 1 if gene family *k* contains at least one gene copy mapped to *a* and one gene copy mapped to *b*, and to 0 otherwise. We define *D*_*MiniNJ*_ :

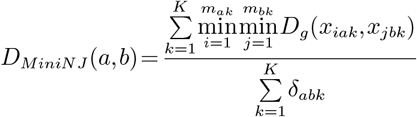

Note that for any two species *a* and *b*, all gene families that cover *a* and *b* contribute equally to *D*_*MiniNJ*_ (*a,b*).

MiniNJ then infers an unrooted species tree from this distance matrix using the NJ algorithm (Saitou and Nei, 1987). The distance matrix computation has time complexity 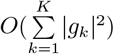 where |*g*_*k*_| is the number of gene sequences in the family *k*. The NJ algorithm has time complexity *O*(|*S*|^3^) where |*S*| is the number of species. The overall time complexity of MiniNJ is thus 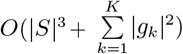.

### Maximum Likelihood rooted species tree search

Given a set *𝒢* of unrooted GFTs, SpeciesRax implements a hill-climbing algorithm to search for the rooted species tree *S* and optimize the set of model parameters Θ (duplication, loss and HGT intensities) that maximize the reconciliation likelihood *L*(*S*,Θ|*𝒢*).

The search starts from an initial species tree *S* and a default or user-specified set of initial model parameters Θ_0_. Then, we alternate between optimizing the species tree root position, Θ, and the species tree topology until we cannot find a configuration with a better likelihood. We describe the exact order in which we execute these distinct steps in the supplementary material. We optimize the root position by evaluating the likelihood of the neighbors of the current root and repeat this process until we do not encounter a neighboring root with a higher likelihood. We optimize Θ via a gradient descent approach. To optimize *S*, we alternate between two complementary tree search strategies that both rely on subtree prune and regraft (SPR) moves: the *transfer-guided SPR search* proposes promising SPR moves by extracting information from the best reconciliation between *S* and *𝒢*. The *local SPR search* tries all possible SPR moves within a user-specified radius (1 by default). In both search strategies, when SpeciesRax finds a species tree *S ′* with a better likelihood than *S*, it replaces *S* by *S ′*. We describe these search operations in more detail in the supplement.

When applying the final root position search, SpeciesRax outputs the per-GFT likelihood scores for all tested root positions. The file with these per-GFT likelihoods can then be further analyzed with the Consel tool (Shimodaira and Hasegawa, 2001) to perform a plethora of statistical significance tests (e.g., the Approximately Unbiased (AU) test (Shimodaira, 2002)) to generate a confident set of root placements.

Calculating the reconciliation likelihood under the UndatedDTL model represents the major computational bottleneck. To reduce its computational cost, we introduce several approximations that we describe in the supplement.

### Support values estimation

Here, we describe how SpeciesRax calculates branch support values on the species tree from a set of unrooted GFTs *𝒢*. We first revisit the definition of a speciation-driven quartet (SQ). Then, we explain how we use the SQ frequency to estimate branch support values. Finally, we describe two alternative SQ-based scores, namely the QPIC and the EQPIC scores.

We first briefly revisit the definition of a SQ (Zhang *et al*., 2019). Let 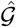 be a set of rooted GFTs with internal nodes either tagged by “duplication” or “speciation” events as estimated from *𝒢*. A quartet from 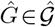 only contains information about the speciation events, if it includes four distinct species *and* if the lowest common ancestor (LCA) of any three out of the four taxa of this quartet is a speciation node. Such a quartet is called SQ. We refer to (Zhang *et al*., 2019) for a more formal definition of the SQ count and for its computation from a set of unrooted and unlabelled GFTs.

We now introduce several notations in order to define the *SQ frequency* of a pair of internal nodes in the species tree. Let *S* be an unrooted species tree. Let (*u,v*) be a pair of distinct internal nodes in *S*. The nodes *u* and *v* define a *metaquartet M*_*u,v*_ =(*A,B,C,D*), where *A* and *B* (resp. *C* and *D*) are the sets of leaves under the left and right children of *u* (resp. *v*) with *S* rooted at *v* (resp. *u*). Let *z* =(*z*_1_,*z*_2_,*z*_3_) such that *z*_1_ (resp. *z*_2_ and *z*_3_) is the *SQ count* in *𝒢* corresponding to the metaquartet topology *AB*|*CD* (resp. *AC*|*BD* and *AD*|*BC*). Note that *z*_1_ corresponds to the metaquartet topology that agrees with *S* ((*A,B*|*C,D*)) and that *z*_2_ and *z*_3_ correspond to the two possible alternative distinct metaquartet topologies (*AC*|*BD* and *AD*|*BC*). Let 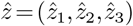 such that 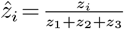 for *i* ∈ (1,2,3). We define the *SQ frequency* of (*u,v*) in *S* given *𝒢* as 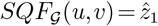.

The SQ frequency represents how many SQs around *u* and *v* support the species tree topology.

However, it does not always reflect if (*AB*|*CD*) is the best supported of the three possible metaquartet topologies, in particular when 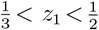 For instance, 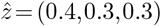 suggests that (*AB*|*CD*) is the correct topology, but 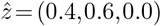 suggests that the alternative topology (*AC*|*BD*) is better supported. Thus, the value of *z*_1_ alone is not sufficiently informative to assess our confidence in a branch.

To overcome this limitation, we therefore also compute the quadripartition internode certainty (QPIC) and extended quadripartition internode certainty (EQPIC) scores introduced in (Zhou *et al*., 2019). Note that these scores were initially defined for single-copy gene families. Since SpeciesRax operates on multiple-copy families, we adapt the scores by only counting SQs instead of counting *all* quartets. Let (*u,v*) be two distinct nodes of *S*.

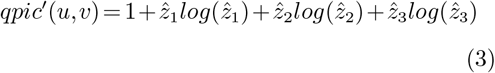

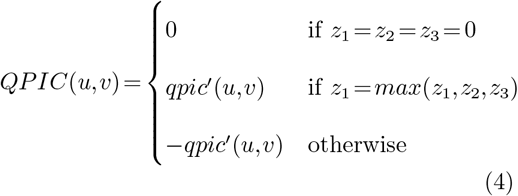

In particular, if *u* and *v* are neighbors, we define the QPIC of the branch *e* between *u* and *v* as *QP IC*(*e*) = *QP IC*(*u,v*). One limitation of the QPIC score is that it discards all SQs defined by nodes *u* and *v* that are not neighbors. (Zhou *et al*., 2019) extends the QPIC score by defining the EQPIC score of a branch *e*:

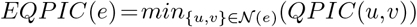

where *N* (*e*) is the set of node pairs *{u,v}* such that the branch *e* belongs to the unique path between *u* and *v*.

We remark that both QPIC and EQPIC scores range between −1 and 1. They take positive values when they support the relevant metaquartet topologies of the species tree *S* and negative values otherwise.

### Accounting for missing data

We refer to *missing data* as gene copies that are absent from a gene family to which they should belong. This can occur, for instance, when some gene sequences have not been sampled or when the gene family clustering is inaccurate. Missing data is problematic for species tree estimation, in particular when the missing data pattern distribution is non-random (Xi *et al*., 2015). In particular, reconciliation methods like SpeciesRax can be affected by missing gene copies: for instance, if sequences for a subset of the species under study have not been sampled for several families, the statistical reconciliation model will attempt to explain these missing gene copies via additional, yet incorrect extinction events. Thus, a candidate species tree that groups such a subset of species into one subtree will typically exhibit a better reconciliation likelihood score than the “true” species tree. This is the case, because only one loss event per family would be necessary to explain all missing gene copies. We alleviate this problem to a certain extent by deploying a *species tree pruning mode*: let *G* be a GFT and *S* a species tree. We replace the reconciliation likelihood term *L*(*S,G*) by *L*(*S′,G*), where *S′* is obtained from *S* by pruning all species that are not covered by *G and* by removing internal nodes of degree 1 until the tree is bifurcating. Thus, if a species is not present in a family, the reconciliation likelihood of this family does not depend on the position of this species in the species tree.

A downside of this approach is that it can disregard some true gene loss events. On both empirical and simulated experiments, we observed that this does not seem to negatively affect the reconstruction accuracy though.

### Parallelization

We parallelized SpeciesRax with MPI (Message Passing Interface) which allows to execute it using several compute nodes with distributed memory (e.g., compute clusters). We distribute the gene families among the available cores to parallelize the reconciliation likelihood computation.

## Experiments

### Tested tools

In the following we describe the settings we used for executing all tools summarized in Table 1 in our experiments. We ran DupTree, FastMulRFS, and MiniNJ with default parameters. Among the four outputs that FastMulRFS provides, we discarded the outputs that may contain multifurcating trees (“majority” and “strict”). Among the two remaining outputs (“greedy” and “single”), we selected “single” because it performed slightly better in our experiments.

**Table 1.** Software used in our benchmark.

| Method | Type | Infers root | Ref. |
| --- | --- | --- | --- |
| NJst | distance matrix | No | (Liu and Yu, 2011) |
| DupTree | parsimony | Yes | (Wehe *et al.*, 2008) |
| FastMulRFS | distance to GFTs | No | (Molloy and Warnow, 2020) |
| ASTRAL-Pro | quartet | No | (Zhang *et al.*, 2019) |
| MiniNJ | distance matrix | No | (this paper) |
| SpeciesRax | maximum likelihood | Yes | (this paper) |

We used our own (re-)implementation of NJst (available in GeneRax) because the existing implementation written in R was too slow for completing our tests in a reasonable time.

We executed ASTRAL-Pro using all available memory (“-Xms700G -Xmx700G”) and a fixed seed (“– seed 692”).

We executed SpeciesRax starting from a MiniNJ tree, with the UndatedDTL model, with per-family duplication, transfer and loss (DTL) rates. We also disabled all irrelevant steps such as gene tree optimization (“-s MiniNJ –optimize-species-tree –do-not-optimize-gene-trees –rec-model UndatedDTL –per-family-rates –skip-family-filtering –do-not-reconcile”). For the experiments on empirical datasets, we added the SpeciesRax option “–prune-species-tree” described in Section to account for missing data. To analyze the empirical dataset that do not contain any multiple-copy gene families (Archaea364), we disabled the gene duplication events in the UndatedDTL model (option “–no-dup”).

### Hardware environment

We executed all experiments on the same machine with 40 physical cores, 80 virtual cores and 750GB RAM. Note that DupTree, FastMulRFS, NJst, and MiniNJ only offer a sequential implementation. In contrast, SpeciesRax and ASTRAL-Pro provide a parallel implementation and were run using all available cores. We discuss the implications of this choice in the results section.

### Simulated datasets

We generated simulated datasets with SimPhy (Mallo *et al*., 2015) to assess the influence of the simulation parameters on the reconstruction accuracy of the methods.

The parameters we studied are: the average number of sites per gene family MSA, the number of families, the size of the species tree, the average DTL rates and the population size. For each parameter we studied, we varied its value while keeping all other parameters fixed. We generated 100 replicates for each set of parameter values. We executed the entire experiment twice, once including HGTs (D**T**LSIM experiment) and once excluding HGTs (DLSIM experiment).

We reused the default parameters of the S25 experiment of (Zhang *et al*., 2019) with some modifications that we list in the following. By default, we do not simulate ILS, which yields the species tree inference easier than in the original S25 experiment. To make the reconstruction more challenging and to reduce the computational cost of the entire experiment, we reduced the number of families from 1000 to 100. To increase the heterogeneity among gene families, we used a log-normal distribution for the sequence length and the DTL rates. In the D**T**LSIM experiment, we simulated under the distance-independent HGT model (i.e., the receiving species is uniformly sampled from all contemporary species) and we set the HGT rates equal to the duplication rates. We provide a detailed list of the SimPhy parameters and arguments used for the gene event rates in the supplement.

We inferred the GFTs with ParGenes (Morel *et al*., 2018), performing one RAxML-NG search on a single random starting tree per gene family under the general time reversible model of nucleotide substitution with four discrete gamma rates (GTR+G) (Tavaré *et al*., 1986; Yang, 1993). Then, we inferred the species trees from the inferred GFTs with every tool listed in Table 1. Finally, for each dataset, we assessed the species tree reconstruction accuracy by computing the average relative RF distance between each inferred species tree and the true species tree using the ETE Toolkit (Huerta-Cepas *et al*., 2016).

### Empirical datasets

We used empirical datasets from various sources to cover a wide range of organisms including plants, fungi, vertebrates, bacteria, and archaea. We describe these datasets in Table. 2. When the datasets included outgroups, we excluded them from the analysis, because SpeciesRax does not need any outgroup to root the species trees. For datasets where we pruned outgroups and for which alignments were available, we reinferred the GFTs from the alignments. This was done to avoid any potential bias in the tree reconstruction that could be caused by the outgroup (Holland *et al*., 2003). In the following we describe in detail how we assembled each empirical dataset.

#### Primates13 and Vertebrates188 datasets

We extracted the alignments comprising 199 species from the Ensembl Compara database. We removed 5 non-vertebrates species to obtain the Vertebrates188 dataset. Further, we extracted 13 primate species to obtain the Primates13 dataset. For both datasets, we inferred the GFTs with ParGenes under the GTR+G model with one random starting tree per RAxML-NG search.

### Cyanobacteria36 dataset

We reused the alignments of a previous study (Szöllősi *et al*., 2013) covering 36 cyanobacteria species to generate the Cyanobacteria36 dataset. We inferred the GFTs with ParGenes under the same substitution model used in the original study (LG+G+I) with one random starting tree per RAxML-NG search.

### Fungi16 and Plants83 datasets

The Fungi16 and Plants83 datasets respectively correspond to the Plant (1kp) and Fungal datasets studied in (Zhang *et al*., 2019). We downloaded the respective GFTs from https://github.com/chaoszhang/A-pro_data.

Fungi60, Plants23 and vertebrates22 datasets We extracted datasets from three different phylomes of the PhylomeDB (Huerta-Cepas *et al*., 2014) database: vertebrates (phylome ID = 200), fungi (phylome ID = 3), and plants (phylome ID = 84). We removed the two outgroup species (*Arabidopsis thaliana* and Human) from the fungi phylome to generate the Fungi60 dataset. We removed the five outgroup species (outgroups: human, Drosophilia, *Caenorhabditis elegans, Saccharomyces cerevisiae* and *Plasmodium falciparum*) from the plant phylome to generate the plants21 dataset. We reinferred the GFTs of both, the fungi and plants datasets using ParGenes with best-fit model selection enabled (-m option) and one random starting tree per RAxML-NG search. We generated the vertebrates22 dataset from the vertebrates phylome without. Here we did not remove any outgroup and did therefore not re-infer the corresponding GFTs.

### Life92 dataset

To compare to the supertrees inferred in the original study (Williams *et al*., 2020), We extracted the original GFTs covering 92 species from the Eukaryote and Archaea domains. To take advantage of the signal from duplications and transfers, we also inferred new homologous gene families from the genomes used in that study. To do so, we performed all-versus-all Diamond (Buchfink *et al*., 2014) searches, then clustered gene families using mcl (Enright, 2002) with an inflation parameter value of 1.4. As in the original study, sequences were aligned using MAFFT (Katoh and Standley, 2013) and poorly-aligning positions removed using BMGE 1.12 (Criscuolo and Gribaldo, 2010) with the BLOSUM30 matrix.

### Archaea364 dataset

We downloaded the MSAs of the marker proteins from the original study (Dombrowski *et al*., 2020). We inferred the GFTss with ParGenes using the LG+G subsitution model.

## Results

### Accuracy on Simphy simulations

We summarize the accuracy of the different species tree reconstruction methods on the D**T**LSIM and DLSIM experiments in Table 2 and Table 3, respectively. We excluded DupTree and NJst from the D**T**LSIM plots and NJst from the DLSIM plots for the sake of an improved visual representation of the results because of their very high error rate.

**Table 2.**
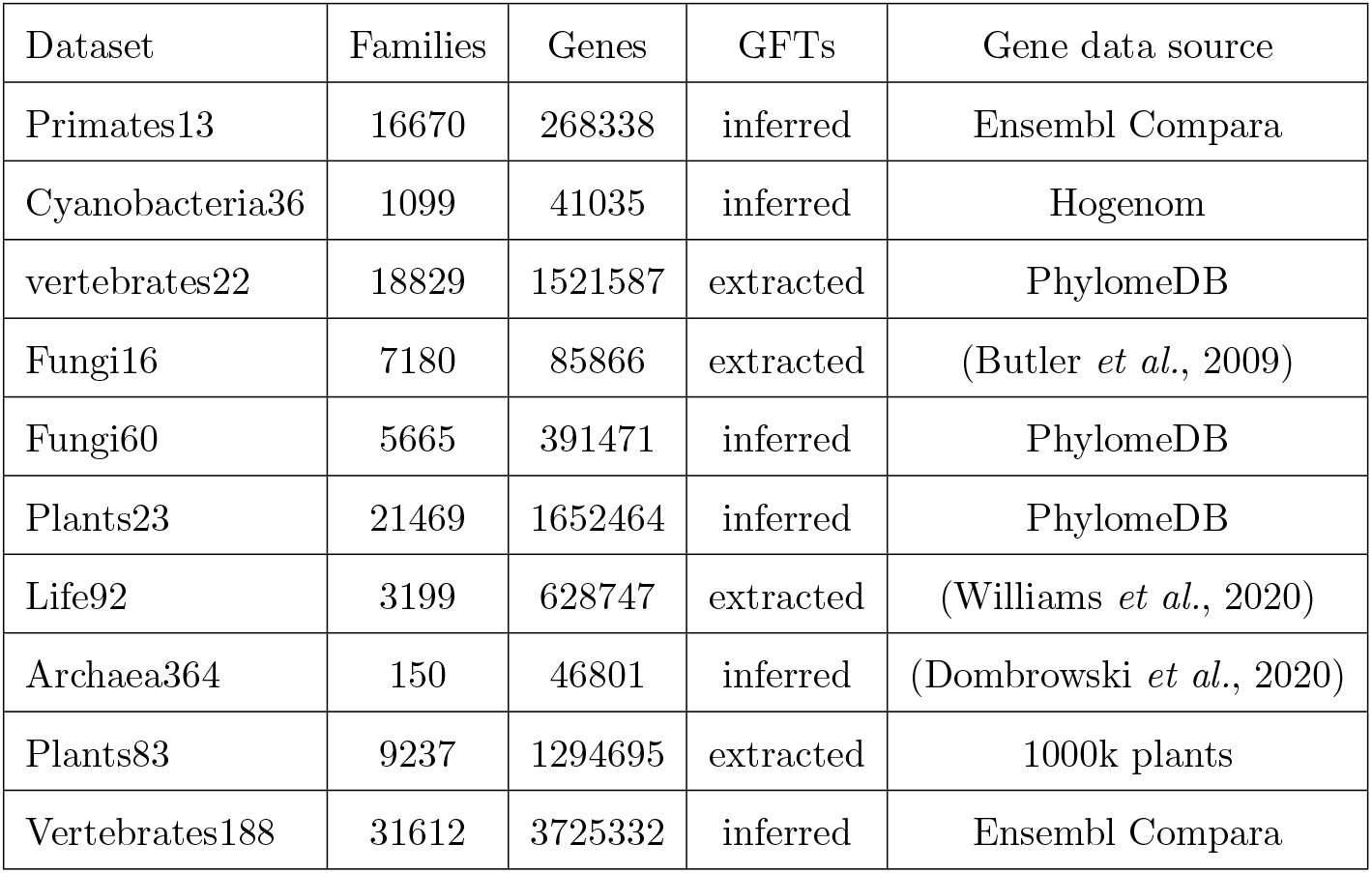
Description of the empirical datasets used in our benchmark. Dataset names are suffixed by the number of species in the respective dataset. Families is the number of input gene families. Genes is the total number of gene copies in the dataset. GFTs indicates if we inferred the GFTs (“inferred”) or if we extracted them from the data source (“extracted”). Gene data source is the database or the project/publication from which the GFTs and/or gene family alignments were obtained.

**Table 3.** Species tree inference runtimes for all tested tools.

| Dataset | FastMulRFS | DupTree | ASTRAL-Pro | SpeciesRax | MiniNJ |
| --- | --- | --- | --- | --- | --- |
| Primates13 | 62s | 18s | 38s | 14s | 2s |
| Cyanobacteria36 | 19s | 26s | 12s | 14s | < 1s |
| Fungi16 | 9s | 7s | 18s | 7s | < 1s |
| Vertebrates22 | 9min | 5min | 2min 45s | 2min 30s | 7s |
| Fungi60 | 16min | 17min | 1min 30s | 1min | 2s |
| Plant23 | 9min | 6min | 1min 35s | 2min | 8s |
| Life92 | 2min 30s | 5min | 1min 30s | 2min | < 1s |
| Archaea364 | 11min | 6h | 1min | 14min | 10s |
| Plants83 | 4h | 2h 40min | 1h 40min | 8min | 27s |
| Vertebrates188 | 14 days | 3.5 days | 12h | 1h 5min | 53s |

In presence of HGTs (D**T**LSIM experiment), SpeciesRax performs better than the competing methods with an average relative RF distance of 0.082 (0.092 for MiniNJ, 0.115 for ASTRAL-Pro, 0.143 for FastMulRFS, 0.409 for NJst and 0.447 for DupTree). In absence of HGTs (DLSIM experiment), we observe the same trend (0.059 for SpeciesRax, 0.063 for MiniNJ, 0.072 for ASTRAL-Pro, 0.089 for FastMulRFS, 0.289 for NJst and 0.116 for DupTree).

As expected, all methods perform better when the phylogenetic signal (number of sites, number of families) increases and perform worse when the discordance between the GFTs and the species tree (ILS level, DTL rates) increases. We do not observe a clear correlation between the number of species and the reconstruction accuracy.

Compared to the other methods, SpeciesRax reconstruction accuracy seems to be less affected by increasing DTL rates and almost unaffected by increasing DL rates. We hypothesize that larger DTL rates increase the species tree - gene trees discordance but also the signal as we obtain larger gene families. Therefore, the competing methods fail to exploit the putative increase in signal but are affected by the higher level of discordance.

Although SpeciesRax does not model ILS, its accuracy is not hampered to a larger degree by increasing population size than that of competing tools.

### Accuracy on empirical datasets

Here, we describe the results of species tree inferences on empirical datasets with ASTRAL-Pro, DupTree, FastMulRFS, MiniNJ, and SpeciesRax. We excluded NJst from this analysis because it performed poorly on most empirical datasets. Initially, we only compare *unrooted* topologies and defer the root placement analysis to a separate subsection. We provide the relative pairwise RF distances between all pairs of inferred trees in the supplement.

#### Vertebrates188 dataset

All tested methods inferred a different species tree. We first counted the number of splits that differ between the inferred trees and the multifurcating NCBI taxonomy (Federhen, 2012) tree. The SpeciesRax, ASTRAL-Pro, and FastMulRFS tools disagree on 5 splits, MiniNJ on 6 splits, and DupTree disagrees on 20 splits.

Then, we focused on the five splits on which SpeciesRax disagrees with the NCBI taxonomy tree that we downloaded from the Ensembl Compara database. Among those discordant splits, the SpeciesRax tree seems to clearly violate only one well established phylogenetic relationship: the elephant shark is believed to have diverged before the split between *Actinopterygii* and *Sarcopterygii* (Venkatesh *et al*., 2014), but SpeciesRax places it as sister to *Sarcopterygii*. Note that all tested methods (ASTRAL-Pro, FastMulRFS, DupTree and MiniNJ) agree with SpeciesRax.

In the following we analyze the remaining four disagreements.

First, SpeciesRax (as well as all other competing tools) places *Cichliformes* as sister to *Ambassidae* while the taxonomy places *Pomacentridae* as sister to *Ambassidae*. Most studies we have found support the taxonomy (Betancur-R *et al*., 2013;

Hughes *et al*., 2018; Near *et al*., 2013)) but other studies are undecided about the resolution of these clades and present trees inferred with different methods that support the three alternative resolutions (Eytan *et al*., 2015).

Another discordance with the taxonomy occurs within the avian subtree, between the *Estrildidae, Fringillidae*, and *Passerellidae* clades: the taxonomy groups the *Estrildidae* and *Fringillidae* together, while SpeciesRax, ASTRAL-Pro, and FastMulRFS group Fringillidae and Passerellidae together. A recently published 363 taxon bird phylogeny (Feng *et al*., 2020) agrees with SpeciesRax on this split and perfectly matches the remaining 24 taxon avian subtree we inferred. In addition, all tested tools place *Bos mutus* (yack) and *Bison bison* closer to each other than to *Bos taurus*, while the taxonomy places *Bos mutus* next to *Bos taurus*. To our knowledge, the literature agrees with our resolution (Decker *et al*., 2009; Kumar *et al*., 2018).

The last inconsistency between the taxonomy and the SpeciesRax tree occurs among the *Platyrrhini* (monkey suborder) when placing *Aotidae, Cebus/Saimiri*, and *Callitrichidae*. This split is perhaps more interesting because SpeciesRax disagrees with the competing methods: the taxonomy places *Cebus/Saimiri* and *Callitrichidae* together, SpeciesRax places *Aotidate* and *Callitrichidae* together. The ASTRAL-Pro, FastMulRFS, MiniNJ, and DupTree tools all group *Aotidate* with *Cebus/Saimiri*. There exist studies that agree with the SpeciesRax (Perelman *et al*., 2011; Springer *et al*., 2012) and the ASTRAL-Pro (Fabre *et al*., 2009) resolutions of these clades.

#### Plants23 dataset

Both SpeciesRax and ASTRAL-Pro species trees disagree with the literature by placing the *Malvales* as sister to *Malpighiales* (instead of sister to *Brassicales* (and V. A. Albert *et al*., 2013; Garcia-Mas *et al*., 2012)). The SpeciesRax species-driven quartet support scores, positively support our resolution, suggesting a potentially misleading signal from the GFTs. When investigating the GFTs, we observed that the *Brassicales* genes often diverge much earlier than they should and that they are often placed outside of the *Rosids* clade to which they should belong. A hypothesis for this misleading signal is the apparent overestimation of the gene family sizes during the gene family clustering performed in the original study (Garcia-Mas *et al*., 2012) as many gene families contain 150 genes (the maximum family size cutoff used in the respective gene family clustering procedure). In addition, the GFTs exhibit clear clusters of genes covering all species separated by extremely long branches. We note however that DupTree and FastMulRFS correctly inferred the entire species tree.

#### Plants83 dataset

The unrooted topologies of the SpeciesRax, ASTRAL-Pro, and FastMulRFS trees are in very 16 good agreement with current biological opinion on the *Viridiplantae* phylogeny, recovering *Setaphyta* and the monophyly of *bryophytes* (Harris *et al*., 2020; Leebens-Mack *et al*., 2019; Puttick *et al*., 2018). The SpeciesRax tree further agrees with several recent analyses (Harris *et al*., 2020; Leebens-Mack *et al*., 2019) in placing the *Coleochaetales* algae as the closest relatives of *Zygnematophyceae* and *Embryophyta* (land plants). The DupTree tree violates many well-established phylogenetic relationships.

#### Fungi60 dataset

All tools found a species tree that disagrees with the literature: they placed the clade formed by *Chytridiomycota* and *Zygomycota* between *Basidiomycota* and *Ascomycota*, which are typically grouped together (Lutzoni *et al*., 2004; Marcet-Houben and Gabaldón, 2009). The positive EQPIC score computed with SpeciesRax along the relevant path shows that the quartets of the GFTs do support this incorrect split. We conclude that the GFTs contain a misleading signal around this split. One possible explanation is that *Encephalitozoon cuniculi* is evolutionary very distant from the remaining species, potentially causing a long branch attraction effect. Apart from this split, SpeciesRax, ASTRAL-Pro, and FastMulRFS inferred the same tree, which agrees with the original species tree obtained via concatenation (Marcet-Houben and Gabaldón, 2009). The tree inferred with DupTree differs from the SpeciesRax tree in one split.

#### Primates13, Cyanobacteria36, Vertebrates22 and Fungi16 datasets

All tools inferred the same species trees for the Primates13, Cyanobacteria36, and Fungi16 datasets and do not violate any well-established phylogenetic relationship. On the Vertebrates22 dataset, all tested methods inferred trees that agree with the multifurcating NCBI taxonomy, but the inferred bifurcating trees are nonetheless different: ASTRAL-Pro, MiniNJ, and SpeciesRax inferred the same tree, which differs from the FastMulRFS tree by one split and the DupTree tree by two splits.

#### Archaea364

The original authors (Dombrowski *et al*., 2020) suggested that one reason for the difficulty in resolving the archaeal tree was the presence of host-symbiont gene transfers in broadly-conserved marker genes, in which members of the DPANN Archaea sometimes grouped with their hosts in single gene phylogenies. Using the full set of marker genes, the SpeciesRax tree recovered a clan (Wilkinson *et al*., 2007) of DPANN; that is, all DPANN Archaea clustered together on the tree. The unrooted SpeciesRax topology is congruent with several recent analyses of the archaeal tree (Dombrowski *et al*., 2020; Raymann *et al*., 2015; Williams *et al*., 2017).

#### Life92

SpeciesRax and ASTRAL-Pro both recovered the major lineages of Archaea and Eukaryotes, including the *Euryarchaeota* and “TACK” Archaea (*Thaumarchaeota, Aigarchaeota, Crenarchaeota* and *Korarchaeota*) within the Archaea, and the SAR, *Archaeplastida* and *Amorphea* clades of Eukaryotes. ASTRAL-Pro resolves the *Excavates* as two separate clades (*Discobans* and *Metamonads*, with *Trimastix* branching between them), while SpeciesRax unites them as sister groups, albeit with very weak statistical support (−0.03); previous work is equivocal as to whether these two lineages form a monophyletic *Excavata* clade (Burki *et al*., 2020; Hampl *et al*., 2009).

SpeciesRax recovers the monophyly of *Asgardarchaeota*, while ASTRAL-Pro instead places one lineage, *Odinarchaeota*, with the TACK Archaea; the position recovered by SpeciesRax is the consensus view (Zaremba-Niedzwiedzka *et al*., 2017). However, SpeciesRax recovers *Asgardarchaeota* as sister to the TACK Archaea, albeit with low support (−0.0075). This topology is incompatible with a specific relationship between Eukaryotes and *Asgardarchaeota*, as supported by analyses of conserved marker genes (Spang *et al*., 2015; Williams *et al*., 2020; Zaremba-Niedzwiedzka *et al*., 2017). The unrooted tree inferred by ASTRAL-Pro groups *Asgardarchaeota* (without *Odinarchaeota*) with Eukaryotes, and is therefore compatible with an origin of the eukaryotic host cell from within the *Asgardarchaeota*.

### Rootings

We now conduct an in depth assessment of the accuracy of the species tree root inference with SpeciesRax on the tested empirical datasets.

We first discuss the datasets on which SpeciesRax inferred a species tree root that agrees with the current literature. On the primates13 dataset, SpeciesRax correctly places the species tree root between the *Strepsirrhini* and *Haplorhini* clades (Chatterjee *et al*., 2009). On the fungi16 dataset, the root inferred with SpeciesRax correctly separates the *Candida* and *Saccharomyces* clades (Butler *et al*., 2009). The root we inferred on the Vertebrates22 species tree correctly separates the *Actinopterygii* and *Sarcopterygii* clades (Meyer and Zardoya, 2003). On the plants23 dataset, our species tree root correctly separates the *Chlorophyta* and *Streptophyta* clades (Leliaert *et al*., 2012). On the fungi60 dataset, we correctly find that *Encephalitozoon cuniculi* (*Microsporidia* clade) diverged earlier than the other clades contained in the dataset (Nagy and Szöllősi, 2017). On the vertebrates188 dataset, SpeciesRax infers a root that groups lampreys and hagfishes, on one side, and cartilaginous fishes, bony fishes, and tetrapodes on the other side. The position of the vertebrate root is still controversial (Miyashita *et al*., 2019; Takezaki *et al*., 2003) and our resolution complies with some of the plausible scenarios discussed in the literature (Meyer and Zardoya, 2003; Miyashita *et al*., 2019; Takezaki *et al*., 2003).

On the plants89 dataset, SpeciesRax agrees with the literature in placing *Embryophyta* (land plants) within the *Streptophyte* algae. However, the inferred root is three branches away from the consensus position, in the common ancestor of the *Chlorophyta* (*Volvox, Chlamydomonas* and *Uronema*).

On the Cyanobacteria36 dataset, the root placement inferred by SpeciesRax is one branch away from one of the three plausible roots inferred in a recent study (Szöllősi *et al*., 2012).

The Archaea364 dataset only contained single-copy gene families, and thus no gene duplications. As a result, the position of the root was uncertain. However the 95% confidence set of possible root placements obtained via the AU test (Shimodaira, 2002) was compatible with several recent suggestions in the literature, including a root between DPANN and all other Archaea (Dombrowski *et al*., 2020; Williams *et al*., 2017) and a root within the Euryarchaeota (Raymann *et al*., 2015), among a range of other positions within and between the major archaeal lineages.

The root inferred by SpeciesRax on the Life92 dataset is biologically not plausible as it is located between *Viridiplantae* and all other taxa. One possibility is that root inference for these data is affected by large differences in gene content among the included taxa. For example, the *Viridiplantae* (and other *Archaeplastida*) have chloroplasts, and so possess an additional source of bacterial-origin genes compared to other Eukaryotes and Archaea. To evaluate the impact of major gene content differences, we performed another SpeciesRax analysis in which the gene families covering less than half of the species were removed. In this second analysis, the root was inferred to lie between the Eukaryotes and Archaea. This root position is compatible with a three-domains tree of life hypothesis. However, this should be interpreted with caution, because the branch separating Eukaryotes and Archaea is one along which major gene content changes occurred, including (but not limited to) the acquisition of a bacterial genome’s worth of genes in the form of the mitochondrial endosymbiont (Roger *et al*., 2017).

### Runtime

Before comparing runtimes, we emphasize again that we executed the experiments on a 40 core machine and that only SpeciesRax and ASTRAL-Pro provide a parallel implementation. While this choice might appear to favor SpeciesRax and ASTRAL-Pro, we argue that the absence of parallelization constitutes a substantial limitation of the remaining tools as completing an analysis in less than one day on a parallel system instead of having to wait for several weeks represents a strong advantage.

We also emphasize that SpeciesRax is the *only* tested tool that can be executed across several compute nodes with distributed memory in contrast to ASTRAL-Pro that can only run on a single shared memory node. All tools, with the exception of MiniNJ, required huge amounts of memory for the largest dataset (*>* 200*GB* on vertebrates188) and can therefore not be executed on most common servers. The SpeciesRax MPI implementation allows to distribute the memory footprint over different compute nodes, which is not feasible with the other tools.

We show the runtimes for an increasing number of species and an increasing number of families for the simulated datasets in Fig. 4. Our MiniNJ method requires less than 0.1 second for all parameter combinations and is the fastest method we tested. The runtimes of DupTree and FastMulRFS grow faster with increasing number of gene families, and DupTree runtime quickly raises with the number of species. The SpeciesRax and ASTRAL-Pro runtimes are less affected by these parameters.

**FIG. 3.**
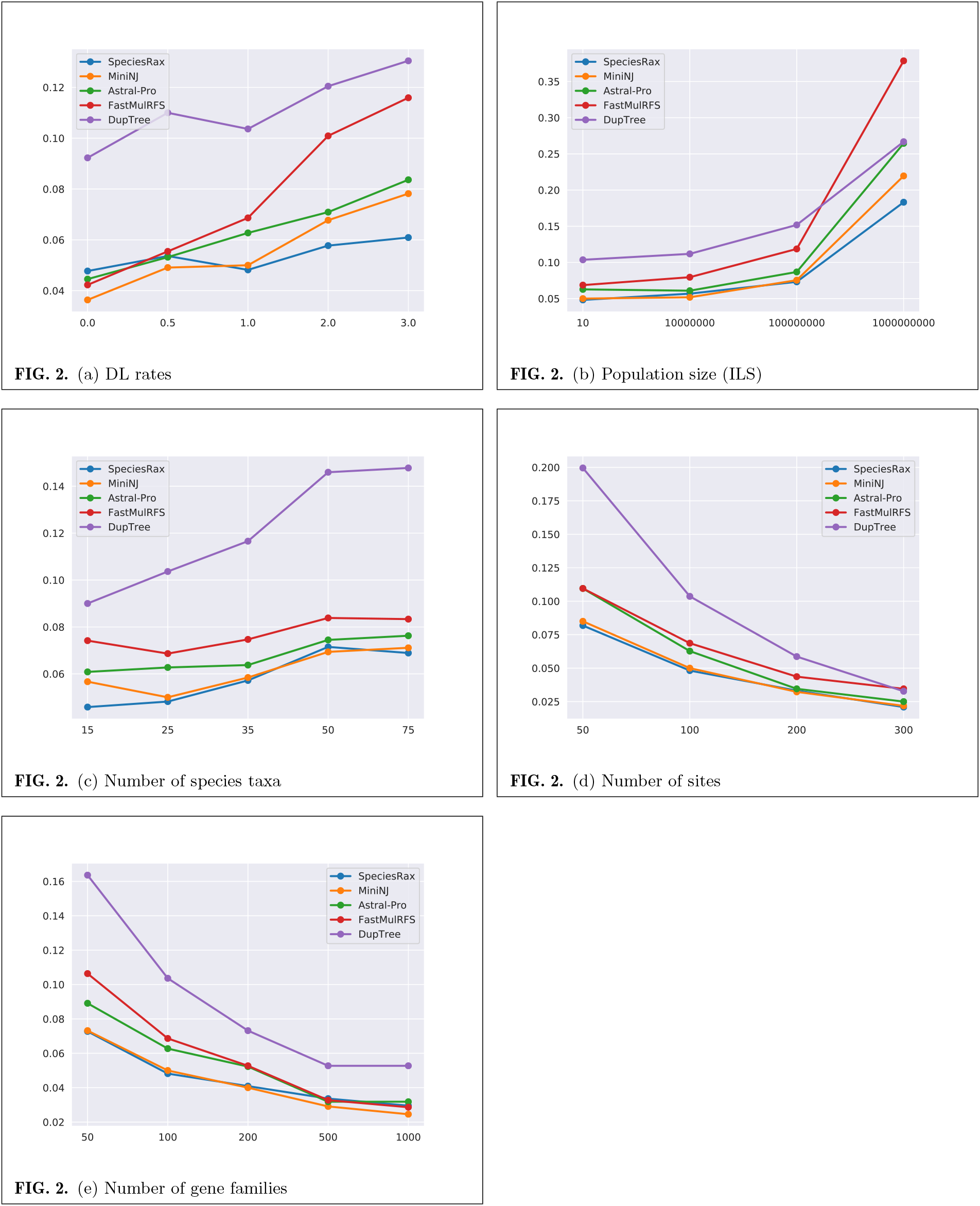
Average unrooted RF distance between inferred and true species trees, in the presence of duplication and loss (no HGTs).

**FIG. 4.**
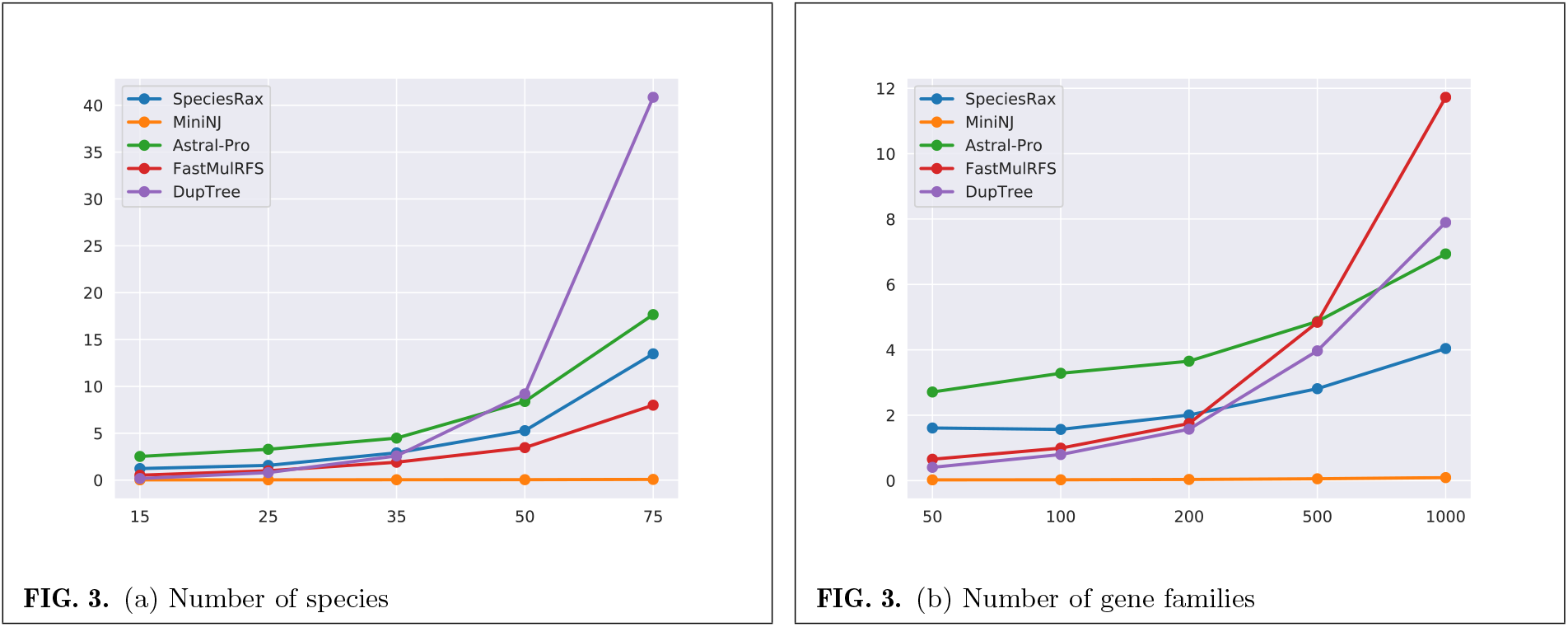
Average runtime in seconds for species tree inference.

On almost all empirical datasets, MiniNJ and SpeciesRax are the fastest methods. On the two largest datasets (Plants83 and Vertebrates 188), MiniNJ is at least one order of magnitude faster than SpeciesRax and SpeciesRax is at least one order of magnitude faster than all other methods. In particular, SpeciesRax only requires one hour on 40 cores to infer the 188 vertebrate species tree with 188 species and 31612 gene families.

## Discussion

### A fast and accurate approach

We introduced two new methods for species tree inference from GFTs in the presence of paralogy. Our MiniNJ tool, is a distance based method that is faster than all tested methods while being at least as accurate as all other non-parametric methods for the majority of our simulated data experiments. In particular, MiniNJ inferred a species tree with 188 species in less than one minute from more than 30000 gene families. SpeciesRax, is a novel maximum likelihood tree search method that explicitly accounts for gene duplication, gene loss, and HGT. Our SpeciesRax tool infers rooted species trees with branch lengths in units of mean expected substitutions per site. Further, to assess the confidence of the inferred species tree, we introduce several quartet based support measures.

In terms of accuracy, SpeciesRax is more accurate than its competitors on simulated datasets, and up to twice as accurate under high duplication, loss, and HGT rates. On empirical datasets, SpeciesRax is on par or more accurate than its competitors. In addition, among the tested tools, SpeciesRax and DupTree are the only methods that can infer *rooted* species trees. SpeciesRax inferred the correct (biologically well-established) roots on 6 out of 10 empirical species trees, and found roots that are close to the plausible roots in 3 out of the remaining 4 datasets (Plants83, Archaea364 and Cyanobacteria). For the most challenging-to-root dataset (Life92), we managed to infer a plausible root by removing those gene families that only covered less than half of the species.

Despite being a compute-intensive maximum likelihood based tree search method, SpeciesRax is faster than all tested methods (except MiniNJ) on large empirical datasets. This is due to our fast method MiniNJ for inferring a reasonable starting tree and to our efficient reconciliation-aware search strategy. In addition, SpeciesRax provides a parallel implementation and can be run on distributed memory cluster systems. Thereby it facilitates conducting large-scale analyses.

Further, SpeciesRax has been integrated into our GeneRax tool that is available via Github and BioConda (Grüning *et al*., 2018). With GeneRax, users can execute the following (optional) steps in one single run: GFT inference from the gene alignments, rooted species tree inference with SpeciesRax from the GFTs, species-tree aware GFT correction, and GFT reconciliation with the rooted species tree. Alternatively, SpeciesRax can be used to infer the root of a user-specified species tree (typically obtained from concatenation methods) before inferring reconciliations. Thus, GeneRax has become a versatile, one-stop shop for executing likelihood based analyses on multiple-copy gene families.

### Future work

Despite our encouraging results, SpeciesRax still faces some challenges.

First, SpeciesRax can currently not take into account GFT reconstruction error/uncertainty. This issue will become more prevalent with increasing taxon numbers and the associated increase in reconstruction uncertainty. Therefore, we intend to explore several ideas to overcome this limitation. A first idea consists in contracting the low-support branches of the GFTs and in adapting our reconciliation model to multifurcating GFTs. Alternatively, we will explore if co-estimating the species tree and the GFTs is feasible, as conducted by Phyldog (Boussau *et al*., 2012), for instance. Finally, we could take as input a distribution of GFTss for each gene family (instead of just one maximum likelihood GFTs) and integrate over this distribution of per gene family GFTs to compute the likelihood score. Such a GFTs distribution could obtained from bayesian inference tools (Ronquist *et al*., 2012), from bootstrap trees (Felsenstein, 1985), or from a set of plausible GFTs (Morel *et al*., 2020).

Finally, we plan to implement more involved models of GFT evolution in SpeciesRax. Some promising work (Chan *et al*., 2017; D Rasmussen and Kellis, 2012; Li *et al*., 2020) has been conducted to account for both DTL events and ILS in a single model. In addition, the UndatedDTL model implemented in SpeciesRax allows for HGTss between any pair of species, even if such HGTss are impossible timewise. Some models (Szöllősi *et al*., 2012) extract time information from the species tree to forbid non-contemporary HGTs, that is, HGTs between species that have not co-existed and thus could not have exchanged genes. We hope that models that better reflect the complexity of gene evolution will yield more reliable species tree inference.

## Data availability

The code is available at https://github.com/BenoitMorel/GeneRax and data are made available at https://cme.h-its.org/exelixis/material/speciesrax_data.tar.gz.

## Acknowledgments

This work was financially supported by the Klaus Tschira Foundation and by DFG grant STA 860/6-2. GSz received funding from the European Research Council under the European Union’s Horizon 2020 research and innovation programme under grant agreement no. 714774 and the grant GINOP-2.3.2.-15-2016-00057. TAW is supported by a Royal Society University Fellowship and NERC grant NE/P00251X/1. This work was funded by the Gordon and Betty Moore Foundation through grant GBMF9741 to TAW and GSz.

## References

Aberer, A.J., Kobert, K., and Stamatakis, A. 2014. ExaBayes: massively parallel bayesian tree inference for the whole-genome era. Molecular Biology and Evolution, 31(10):2553–2556.

Altenhoff, A.M., Glover, N.M., and Dessimoz, C. 2019. Inferring Orthology and Paralogy, pages 149–175. Springer New York, New York, NY.

and V. A. Albert, Barbazuk, W.B., dePamphilis, C.W., Der, J.P., Leebens-Mack, J., Ma, H., Palmer, J.D., Rounsley, S., Sankoff, D., Schuster, S.C., Soltis, D.E., Soltis, P.S., Wessler, S.R., Wing, R.A., Albert, V.A., Ammiraju, J.S. S., Barbazuk, W.B., Chamala, S., Chanderbali, A.S., dePamphilis, C.W., Der, J.P., Determann, R., Leebens-Mack, J., Ma, H., Ralph, P., Rounsley, S., Schuster, S.C., Soltis, D.E., Soltis, P.S., Talag, J., Tomsho, L., Walts, B., Wanke, S., Wing, R.A., Albert, V.A., Barbazuk, W.B., Chamala, S., Chanderbali, A.S., Chang, T.-H., Determann, R., Lan, T., Soltis, D.E., Soltis, P.S., Arikit, S., Axtell, M.J., Ayyampalayam, S., Barbazuk, W.B., Burnette, J.M., Chamala, S., Paoli, E.D., dePamphilis, C.W., Der, J.P., Estill, J.C., Farrell, N.P., Harkess, A., Jiao, Y., Leebens-Mack, J., Liu, K., Mei, W., Meyers, B.C., Shahid, S., Wafula, E., Walts, B., Wessler, S.R., Zhai, J., Zhang, X., Albert, V.A., Carretero-Paulet, L., dePamphilis, C.W., Der, J.P., Jiao, Y., Leebens-Mack, J., Lyons, E., Sankoff, D., Tang, H., Wafula, E., Zheng, C., Albert, V.A., Altman, N.S., Barbazuk, W.B., Carretero-Paulet, L., dePamphilis, C.W., Der, J.P., Estill, J.C., Jiao, Y., Leebens-Mack, J., Liu, K., Mei, W., Wafula, E., Altman, N.S., Arikit, S., Axtell, M.J., Chamala, S., Chanderbali, A.S., Chen, F., Chen, J.-Q., Chiang, V., Paoli, E.D., dePamphilis, C.W., Der, J.P., Determann, R., Fogliani, B., Guo, C., Harholt, J., Harkess, A., Job, C., Job, D., Kim, S., Kong, H., Leebens-Mack, J., Li, G., Li, L., Liu, J., Ma, H., Meyers, B.C., Park, J., Qi, X., Rajjou, L., Burtet-Sarramegna, V., Sederoff, R., Shahid, S., Soltis, D.E., Soltis, P.S., Sun, Y.-H., Ulvskov, P., Villegente, M., Xue, J.-Y., Yeh, T.-F., Yu, X., Zhai, J., Acosta, J.J., Albert, V.A., Barbazuk, W.B., Bruenn, R.A., Chamala, S., de Kochko, A., dePamphilis, C.W., Der, J.P., Herrera-Estrella, L.R., Ibarra-Laclette, E., Kirst, M., Leebens-Mack, J., Pissis, S.P., Poncet, V., Schuster, S.C., Soltis, D.E., Soltis, P.S., and Tomsho, L. 2013. The amborella genome and the evolution of flowering plants. Science, 342(6165):1241089–1241089.

Bayzid, M., Mirarab, S., and Warnow, T. 2013. Inferring optimal species trees under gene duplication and loss. Pacific Symposium on Biocomputing. Pacific Symposium on Biocomputing, pages 250–261. 18th Pacific Symposium on Biocomputing, PSB 2013; Conference date: 03-01-2013 Through 07-01-2013.

Betancur-R, R., Broughton, R.E., Wiley, E.O., Carpenter, K., Lopez, J.A., Li, C., Holcroft, N.I., Arcila, D., Sanciangco, M., Cureton Ii, J.C., et al. 2013. The tree of life and a new classification of bony fishes. PLoS currents, 5.

Bouckaert, R., Heled, J., Kühnert, D., Vaughan, T., Wu, C.-H., Xie, D., Suchard, M.A., Rambaut, A., and Drummond, A.J. 2014. Beast 2: A software platform for bayesian evolutionary analysis. PLOS Computational Biology, 10(4):1–6.

Boussau, B., Szöllősi, G.J., Duret, L., Gouy, M., Tannier, E., Daubin, V., Lyon, U.D., and Lyon, U. 2012. Genome-scale coestimation of species and gene trees. Life Sciences, pages 1–27.

Buchfink, B., Xie, C., and Huson, D.H. 2014. Fast and sensitive protein alignment using DIAMOND. Nature Methods, 12(1):59–60.

Burki, F., Roger, A.J., Brown, M.W., and Simpson, A.G. 2020. The new tree of eukaryotes. Trends in Ecology & Evolution, 35(1):43–55.

Butler, G., Rasmussen, M., Lin, M., Sakthikumar, S., Munro, C., Rheinbay, E., Grabherr, M., Forche, A., Reedy, J., Agrafioti, I., Arnaud, M., Bates, S., Brown, A., Brunke, S., Costanzo, M., Fitzpatrick, D., Groot, P., Harris, D., and Cuomo, C. 2009. Evolution of pathogenicity and sexual reproduction in eight candida genomes. Nature, 459:657–62.

Chan, Y., Ranwez, V., and Scornavacca, C. 2017. Inferring incomplete lineage sorting, duplications, transfers and losses with reconciliations. Journal of Theoretical Biology, 432:1 – 13.

Chatterjee, H.J., Ho, S.Y., Barnes, I., and Groves, C. 2009. Estimating the phylogeny and divergence times of primates using a supermatrix approach. BMC Evolutionary Biology, 9(1):259.

Criscuolo, A. and Gribaldo, S. 2010. BMGE (block mapping and gathering with entropy): a new software for selection of phylogenetic informative regions from multiple sequence alignments. BMC Evolutionary Biology, 10(1):210.

D Rasmussen, M. and Kellis, M. 2012. Unified modeling of gene duplication, loss, and coalescence using a locus tree. Genome research, 22:755–65.

de Oliveira Martins, L. and Posada, D. 2017. Species Tree Estimation from Genome-Wide Data with guenomu, pages 461–478. Springer New York, New York, NY.

Decker, J.E., Pires, J.C., Conant, G.C., McKay, S.D., Heaton, M.P., Chen, K., Cooper, A., Vilkki, J., Seabury, C.M., Caetano, A.R., Johnson, G.S., Brenneman, B.C., Petersen, B., Wang, Z., Zhou, Q., Diekhans, M., Chen, W., Andreu-Śanchez, S., Margaryan, A., Howard, J.T., Parent, C., Pacheco, G., Sinding, M.H. S., Puetz, R.A., Hanotte, O., Eggert, L.S., Wiener, P., Kim, L., Cavill, E., Ribeiro Â M., Eckhart, L., Fjeldså, J.-J., Kim, K.S., Sonstegard, T.S., Van Tassell, C.P., Neibergs, H.L., McEwan, J.C., Brauning, R., Coutinho, L.L., Babar, M.E., Wilson, G.A., McClure, M.C., Rolf, M.M., Kim, J., Schnabel, R.D., and Taylor, J.F. 2009. Resolving the evolution of extant and extinct ruminants with high-throughput phylogenomics. Proceedings of the National Academy of Sciences, 106(44):18644–18649.

Dombrowski, N., Williams, T.A., Sun, J., Woodcroft, B.J., Lee, J.-H., Minh, B.Q., Rinke, C., and Spang, A. 2020. Undinarchaeota illuminate DPANN phylogeny and the impact of gene transfer on archaeal evolution. Nature Communications, 11(1).

Emms, D. and Kelly, S. 2018. Stag: Species tree inference from all genes. bioRxiv.

Enright, A.J. 2002. An efficient algorithm for large-scale detection of protein families. Nucleic Acids Research, 30(7):1575–1584.

Eytan, R.I., Evans, B.R., Dornburg, A., Lemmon, A.R., Lemmon, E.M., Wainwright, P.C., and Near, T.J. 2015. Are 100 enough? inferring acanthomorph teleost phylogeny using anchored hybrid enrichment. BMC Evolutionary Biology, 15(1).

Fabre, P.-H., Rodrigues, A., and Douzery, E. 2009. Patterns of macroevolution among primates inferred from a supermatrix of mitochondrial and nuclear dna. Molecular Phylogenetics and Evolution, 53(3):808 – 825.

Federhen, S. 2012. The NCBI Taxonomy database. Nucleic Acids Research, 40(D1): 136–143.

Felsenstein, J. 1985. Confidence limits on phylogenies: an approach using the bootstrap. Evolution, 39(4):783– 791.

Feng, S., Stiller, J., Deng, Y., Armstrong, J., Fang, Q., Reeve, A.H., Xie, D., Chen, G., Guo, C., Faircloth, J., Hosner, P.A., Brumfield, R.T., Christidis, L., Bertelsen, M.F., Sicheritz-Ponten, T., Tietze, D.T., Robertson, B.C., Song, G., Borgia, G., Claramunt, S., Lovette, I.J., Cowen, S.J., Njoroge, P., Dumbacher, J.P., Ryder, O.A., Fuchs, J., Bunce, M., Burt, D.W., Cracraft, J., Meng, G., Hackett, S.J., Ryan, P.G., Jønsson, K.A., Jamieson, I.G., da Fonseca, R.R., Braun, E.L., Houde, P., Mirarab, S., Suh, A., Hansson, B., Ponnikas, S., Sigeman, H., Stervander, M., Frandsen, P.B., van der Zwan, H., van der Sluis, R., Visser, C., Balakrishnan, C.N., Clark, A.G., Fitzpatrick, J.W., Bowman, R., Chen, N., Cloutier, A., Sackton, T.B., Edwards, S.V., Foote, D.J., Shakya, S.B., Sheldon, F.H., Vignal, A., Soares, A.E., Shapiro, B., González-Solís, J., Ferrer-Obiol, J., Rozas, J., Riutort, M., Tigano, A., Friesen, V., Dalén, L., Urrutia, A.O., Székely, T., Liu, Y., Campana, M.G., Corvelo, A., Fleischer, R.C., Rutherford, K.M., Gemmell, N.J., Dussex, N., Mouritsen, H., Thiele, N., Delmore, K., Liedvogel, M., Franke, A., Hoeppner, M.P., Krone, O., Fudickar, A.M., Milá, B., Ketterson, E.D., Fidler, A.E., Friis, G., Parody-Merino, Á.M., Battley, P.F., Cox, M.P., Lima, N.C. B., Prosdocimi, F., Parchman, T.L., Schlinger, B.A., Loiselle, B.A., Blake, J.G., Lim, H.C., Day, L.B., Fuxjager, M.J., Baldwin, M.W., Braun, M.J., Wirthlin, M., Dikow, R.B., Ryder, T.B., Camenisch, G., Keller, L.F., DaCosta, J.M., Hauber, M.E., Louder, M.I., Witt, C.C., McGuire, J.A., Mudge, J., Megna, L.C., Carling, M.D., Wang, B., Taylor, S.A., Del-Rio, G., Aleixo, A., Vasconcelos, A.T. R., Mello, C.V., Weir, J.T., Haussler, D., Li, Q., Yang, H., Wang, J., Lei, F., Rahbek, C., Gilbert, M.T. P., Graves, G.R., Jarvis, E.D., Paten, B., and Zhang, G. 2020. Dense sampling of bird diversity increases power of comparative genomics. Nature, 587(7833):252–257.

Garcia-Mas, J., Benjak, A., Sanseverino, W., Bourgeois, M., Mir, G., Gonzalez, V.M., Henaff, E., Camara, F., Cozzuto, L., Lowy, E., Alioto, T., Capella-Gutierrez, S., Blanca, J., Canizares, J., Ziarsolo, P., Gonzalez-Ibeas, D., Rodriguez-Moreno, L., Droege, M., Du, L., Alvarez-Tejado, M., Lorente-Galdos, B., Mele, M., Yang, L., Weng, Y., Navarro, A., Marques-Bonet, T., Aranda, M.A., Nuez, F., Pico, B., Gabaldon, T., Roma, G., Guigo, R., Casacuberta, J.M., Arus, P., and Puigdomenech, P. 2012. The genome of melon (cucumis melo l.). Proceedings of the National Academy of Sciences, 109(29):11872–11877.

Grüning, B., Dale, R., Sjödin, A., Chapman, B., Rowe, J., Tomkins-Tinch, C., Valieris, R., Köster, J., Blin, K., Haudgaard, M., Kratz, A., Junge, A., and Knudsen, M. 2018. Bioconda: sustainable and comprehensive software distribution for the life sciences. Nature Methods, 15:475–476.

Hampl, V., Hug, L., Leigh, J.W., Dacks, J.B., Lang, B.F., Simpson, A.G. B., and Roger, A.J. 2009. Phylogenomic analyses support the monophyly of excavata and resolve relationships among eukaryotic “supergroups”. Proceedings of the National Academy of Sciences, 106(10):3859–3864.

Harris, B.J., Harrison, C.J., Hetherington, A.M., and Williams, T.A. 2020. Phylogenomic evidence for the monophyly of bryophytes and the reductive evolution of stomata. Current Biology, 30(11):2001–2012.e2.

Holland, B.R., Penny, D., and Hendy, M.D. 2003. Outgroup Misplacement and Phylogenetic Inaccuracy Under a Molecular Clock—A Simulation Study. Systematic Biology, 52(2):229–238.

Huerta-Cepas, J., Capella-Gutiérrez, S., Pryszcz, L., Marcet-Houben, M., and Gabaldón, T. 2014. Phylomedb v4: zooming into the plurality of evolutionary histories of a genome. Nucleic Acids Research, 42:D897 – D902.

Huerta-Cepas, J., Serra, F., and Bork, P. 2016. ETE 3: Reconstruction, Analysis, and Visualization of Phylogenomic Data. Molecular Biology and Evolution, 33(6):1635–1638.

Hughes, L.C., Ortí, G., Huang, Y., Sun, Y., Baldwin, C.C., Thompson, A.W., Arcila, D., Betancur-R., R., Li, C., Becker, L., Bellora, N., Zhao, X., Li, X., Wang, M., Fang, C., Xie, B., Zhou, Z., Huang, H., Chen, S., Venkatesh, B., and Shi, Q. 2018. Comprehensive phylogeny of ray-finned fishes (actinopterygii) based on transcriptomic and genomic data. Proceedings of the National Academy of Sciences, 115(24):6249–6254.

Katoh, K. and Standley, D.M. 2013. MAFFT Multiple Sequence Alignment Software Version 7: Improvements in Performance and Usability. Molecular Biology and Evolution, 30(4):772–780.

Kozlov, A.M., Darriba, D., Flouri, T., Morel, B., and Stamatakis, A. 2019. RAxML-NG: a fast, scalable and user-friendly tool for maximum likelihood phylogenetic inference. Bioinformatics, 35(21):4453–4455.

Kubatko, L.S. and Degnan, J.H. 2007. Inconsistency of Phylogenetic Estimates from Concatenated Data under Coalescence. Systematic Biology, 56(1):17–24.

Kumar, P., Velayutham, D.pkS.Ps B., Zachariah, A., Zachariah, A., Bathrachalam, C., Sajeevkumar, S.S.G., Bangarusamy, D., Iype, S., Gupta, R., Santhosh, S., and Thomas, G. 2018. Complete mitogenome reveals genetic divergence and phylogenetic relationships among indian cattle (bos indicus) breeds. Animal Biotechnology, 30:1–14.

Leebens-Mack, J.H., Barker, M.S., Carpenter, E.J., Deyholos, M.K., Gitzendanner, M.A., Graham, S.W., Grosse, I., Li, Z., Melkonian, M., Mirarab, S., et al. 2019. One thousand plant transcriptomes and the phylogenomics of green plants. Nature, 574(7780):679–685.

Leliaert, F., Smith, D.R., Moreau, H., Herron, M.D., Verbruggen, H., Delwiche, C.F., and Clerck, O.D. 2012. Phylogeny and molecular evolution of the green algae. Critical Reviews in Plant Sciences, 31(1):1–46.

Li, Q., Scornavacca, C., Galtier, N., and Chan, Y.-B. 2020. The Multilocus Multispecies Coalescent: A Flexible New Model of Gene Family Evolution. Systematic Biology. syaa084.

Liu, L. and Yu, L. 2011. Estimating Species Trees from Unrooted Gene Trees. Systematic Biology, 60(5):661– 667.

Lutzoni, F., Kauff, F., Cox, C., McLaughlin, D., Celio, G., Dentinger, B., Padamsee, M., Hibbett, D., James, T., Baloch, E., Grube, M., Reeb, V., Valerie, H., Schoch, C., Arnold, A., Miadlikowska, J., Spatafora, J., Johnson, D., Hambleton, S., and Vilgalys, R. 2004. Assembling the fungal tree of life: Progress, classification, and evolution of subcellular traits. American journal of botany, 91:1446–80.

Mallo, D., De Oliveira Martins, L., and Posada, D. 2015. SimPhy : Phylogenomic Simulation of Gene, Locus, and Species Trees. Systematic Biology, 65(2):334–344.

Marcet-Houben, M. and Gabaldon, T. 2009. The Tree versus the forest: The fungal tree of life and the topological diversity within the yeast phylome. PLoS ONE, 4(2).

Mendes, F.K. and Hahn, M.W. 2017. Why Concatenation Fails Near the Anomaly Zone. Systematic Biology, 67(1):158–169.

Meyer, A. and Zardoya, R. 2003. Recent advances in the (molecular) phylogeny of vertebrates. Annual Review of Ecology and Systematics 34 (2003), pp. 311–338, 34.

Minh, B.Q., Schmidt, H.A., Chernomor, O., Schrempf, D., Woodhams, M.D., von Haeseler, A., and Lanfear, R. 2020. IQ-TREE 2: New Models and Efficient Methods for Phylogenetic Inference in the Genomic Era. Molecular Biology and Evolution, 37(5):1530–1534.

Miyashita, T., Coates, M.I., Farrar, R., Larson, P., Manning, P.L., Wogelius, R.A., Edwards, N.P., Anné, J., Bergmann, U., Palmer, A.R., and Currie, P.J. 2019. Hagfish from the cretaceous tethys sea and a reconciliation of the morphological–molecular conflict in early vertebrate phylogeny. Proceedings of the National Academy of Sciences, 116(6):2146–2151.

Molloy, E.K. and Warnow, T. 2020. FastMulRFS: fast and accurate species tree estimation under generic gene duplication and loss models. Bioinformatics, 36(Supplement 1): i57–i65.

Morel, B., Kozlov, A.M., and Stamatakis, A. 2018. ParGenes: a tool for massively parallel model selection and phylogenetic tree inference on thousands of genes. Bioinformatics.

Morel, B., Kozlov, A.M., Stamatakis, A., and Szöllősi, G.J. 2019. Generax: A tool for species tree-aware maximum likelihood based gene tree inference under gene duplication, transfer, and loss. bioRxiv.

Morel, B., Barbera, P., Czech, L., Bettisworth, B., Hübner, L., Lutteropp, S., Serdari, D., Kostaki, E.-G., Mamais, I., Kozlov, A.M., Pavlidis, P., Paraskevis, D., and Stamatakis, A. 2020. Phylogenetic Analysis of SARS-CoV-2 Data Is Difficult. Molecular Biology and Evolution. msaa 314.

Nagy, L.G. and Szöllősi, G. 2017. Chapter two - fungal phylogeny in the age of genomics: Insights into phylogenetic inference from genome-scale datasets. In J. P. Townsend and Z. Wang, editors, Fungal Phylogenetics and Phylogenomics, volume 100 of Advances in Genetics, pages 49–72. Academic Press.

Near, T.J., Dornburg, A., Eytan, R.I., Keck, B.P., Smith, W.L., Kuhn, K.L., Moore, J.A., Price, S.A., Burbrink, F.T., Friedman, M., and Wainwright, P.C. 2013. Phylogeny and tempo of diversification in the superradiation of spiny-rayed fishes. Proceedings of the National Academy of Sciences, 110(31):12738–12743.

Perelman, P., Johnson, W.E., Roos, C., Seuánez, H.N., Horvath, J.E., Moreira, M.A. M., Kessing, B., Pontius, J., Roelke, M., Rumpler, Y., Schneider, M.P. C., Silva, A., O’Brien, S.J., and Pecon-Slattery, J. 2011. A molecular phylogeny of living primates. PLOS Genetics, 7(3):1–17.

Puttick, M.N., Morris, J.L., Williams, T.A., Cox, C.J., Edwards, D., Kenrick, P., Pressel, S., Wellman, C.H., Schneider, H., Pisani, D., and Donoghue, P.C. 2018. The interrelationships of land plants and the nature of the ancestral embryophyte. Current Biology, 28(5):733–745.e2.

Raymann, K., Brochier-Armanet, C., and Gribaldo, S. 2015. The two-domain tree of life is linked to a new root for the archaea. Proceedings of the National Academy of Sciences, 112(21):6670–6675.

Roger, A.J., Munóz-Gómez, S.A., and Kamikawa, R. 2017. The origin and diversification of mitochondria. Current Biology, 27(21):R1177–R1192.

Ronquist, F., Teslenko, M., van der Mark, P., Ayres, D.L., Darling, A., Höhna, S., Larget, B., Liu, L., Suchard, M.A., and Huelsenbeck, J.P. 2012. MrBayes 3.2: Efficient Bayesian Phylogenetic Inference and Model Choice Across a Large Model Space. Systematic Biology, 61(3):539–542.

Saitou, N. and Nei, M. 1987. The neighbor-joining method: a new method for reconstructing phylogenetic trees. Molecular biology and evolution, 4(4):406–425.

Shimodaira, H. 2002. An approximately unbiased test of phylogenetic tree selection. Systematic Biology, 51(3):492–508.

Shimodaira, H. and Hasegawa, M. 2001. CONSEL: for assessing the confidence of phylogenetic tree selection. Bioinformatics, 17(12):1246–1247.

Spang, A., Saw, J.H., Jørgensen, S.L., Zaremba-Niedzwiedzka, K., Martijn, J., Lind, A.E., van Eijk, R., Schleper, C., Guy, L., and Ettema, T.J. G. 2015. Complex archaea that bridge the gap between prokaryotes and eukaryotes. Nature, 521(7551):173– 179.

Springer, M.S., Meredith, R.W., Gatesy, J., Emerling, C.A., Park, J., Rabosky, D.L., Stadler, T., Steiner, C., Ryder, O.A., Janečka, J.E., Fisher, C.A., and Murphy, W.J. 2012. Macroevolutionary dynamics and historical biogeography of primate diversification inferred from a species supermatrix. PLOS ONE, 7(11):1–23.

Szöllősi, G.J., Rosikiewicz, W., Boussau, B., Tannier, E., and Daubin, V. 2013. Efficient exploration of the space of reconciled gene trees. Systematic Biology, 62(6):901– 912.

Szöllősi, G.J., Boussau, B., Abby, S.S., Tannier, E., and Daubin, V. 2012. Phylogenetic modeling of lateral gene transfer reconstructs the pattern and relative timing of speciations. Proceedings of the National Academy of Sciences, 109(43):17513–17518.

Takezaki, N., Figueroa, F., Zaleska-Rutczynska, Z., and Klein, J. 2003. Molecular Phylogeny of Early Vertebrates: Monophyly of the Agnathans as Revealed by Sequences of 35 Genes. Molecular Biology and Evolution, 20(2):287–292.

Tavaré, S. et al.. 1986. Some probabilistic and statistical problems in the analysis of dna sequences. Lectures on mathematics in the life sciences, 17(2):57–86.

Venkatesh, B., Lee, A.P., Ravi, V., Maurya, A.K., Lian, M.M., Swann, J.B., Ohta, Y., Flajnik, M.F., Sutoh, Y., Kasahara, M., Hoon, S., Gangu, V., Roy, S.W., Irimia, M., Korzh, V., Kondrychyn, I., Lim, Z.W., Tay, B.H., Tohari, S., Kong, K.W., Ho, S., Lorente-Galdos, B., Quilez, J., Marques-Bonet, T., Raney, B.J., Ingham, P.W., Tay, A., Hillier, L.W., Minx, P., Boehm, T., Wilson, R.K., Brenner, S., and Warren, W.C. 2014. Elephant shark genome provides unique insights into gnathostome evolution. Nature, 505(7482):174–179.

Wehe, A., Bansal, M.S., Burleigh, J.G., and Eulenstein, O. 2008. DupTree: a program for large-scale phylogenetic analyses using gene tree parsimony. Bioinformatics, 24(13):1540–1541.

Wilkinson, M., McInerney, J.O., Hirt, R.P., Foster, P.G., and Embley, T.M. 2007. Of clades and clans: terms for phylogenetic relationships in unrooted trees. Trends in Ecology & Evolution, 22(3):114–115.

Williams, T.A. and Embley, T.M. 2014. Archaeal “Dark Matter” and the Origin of Eukaryotes. Genome Biology and Evolution, 6(3):474–481.

Williams, T.A., Szöllosi, G.J., Spang, A., Foster, P.G., Heaps, S.E., Boussau, B., Ettema, T.J. G., and Embley, T.M. 2017. Integrative modeling of gene and genome evolution roots the archaeal tree of life. Proceedings of the National Academy of Sciences, 114(23):E4602– E4611.

Williams, T.A., Cox, C.J., Foster, P.G., Szöllősi, G.J., and Embley, T.M. 2020. Phylogenomics provides robust support for a two-domains tree of life. Nature Ecology & Evolution, 4(1):138–147.

Xi, Z., Liu, L., and Davis, C.C. 2015. The Impact of Missing Data on Species Tree Estimation. Molecular Biology and Evolution, 33(3):838–860.

Yang, Z. 1993. Maximum-likelihood estimation of phylogeny from dna sequences when substitution rates differ over sites. Molecular biology and evolution, 10(6):1396–1401.

Zaremba-Niedzwiedzka, K., Caceres, E.F., Saw, J.H., Bäckström, D., Juzokaite, L., Vancaester, E., Seitz, K.W., Anantharaman, K., Starnawski, P., Kjeldsen, K.U., Stott, M.B., Nunoura, T., Banfield, J.F., Schramm, A., Baker, B.J., Spang, A., and Ettema, T.J. G. 2017. Asgard archaea illuminate the origin of eukaryotic cellular complexity. Nature, 541(7637):353–358.

Zhang, C., Sayyari, E., and Mirarab, S. 2017. Astral-iii: Increased scalability and impacts of contracting low support branches. In Comparative Genomics, pages 53–75, Cham. Springer International Publishing.

Zhang, C., Scornavacca, C., Molloy, E.K., and Mirarab, S. 2019. Astral-pro: quartet-based species tree inference despite paralogy. bioRxiv.

Zhou, X., Lutteropp, S., Czech, L., Stamatakis, A., Looz, M.V., and Rokas, A. 2019. Quartet-Based Computations of Internode Certainty Provide Robust Measures of Phylogenetic Incongruence. Systematic Biology, 69(2):308–324.

