## Supplement material for "SpeciesRax: A tool for maximum likelihood species tree inference from gene family trees under duplication, transfer, and loss"

### 1 Reconciliation likelihood

#### 1.1 Exact reconciliation likelihood computation

Let $S$ be a rooted species tree and $G$ be a rooted gene tree. We briefly outline how we compute the reconciliation likelihood $P(G|S)$. A more detailed description is provided in [Morel *et al.*(2019)Morel, Kozlov, Stamatakis, and Szöllősi].

Let $|S|$ and $|G|$ be the number of taxa in $S$ and $G$, respectively. Let $u$ be a branch of $G$ and let $v$ and $w$ be its descendant branches. Let $e$ be a branch of $S$ and let $f$ and $g$ be its descendant branches.

Let $P^S$, $P^D$, $P^L$, and $P^T$ be the speciation, duplication, loss, and transfer probabilities and let $\delta$, $\lambda$, and $\tau$ be the duplication, loss, and transfer intensity parameters that parametrize the duplication event probabilities as follows:

$$p^D = \delta/(1+\delta+\tau+\lambda) \tag{1}$$
$$p^L = \lambda/(1+\delta+\tau+\lambda) \tag{2}$$
$$p^T = \tau/(1+\delta+\tau+\lambda) \tag{3}$$
$$p^S = 1/(1+\delta+\tau+\lambda). \tag{4}$$

The extinction probability, that is, the probability that a gene copy observed on an internal branch $e$ becomes extinct before being observed at the tips of the specie tree is:

$$E_e = p^L + p^S\left(E_f E_g\right) + p^D\left(E_e^2\right) + p^T\left(E_e \bar{E}_e\right). \tag{5}$$

with

$$\bar{E}_e = \sum_{h \in \mathcal{T}(e)} \frac{E_h}{|\mathcal{T}(e)|} \tag{6}$$

Let $P_{e,u}$ be the probability of observing the internal branch $u$ of $G$ on the internal branch $e$ of $S$. We can write $P_{e,u}$ as:

$$\begin{aligned} P_{e,u} = {} & p^S\left(P_{g,v}P_{f,w} + P_{g,w}P_{f,v}\right) + p^S\left(E_f P_{g,u} + P_{f,u}E_g\right) \\ & + p^D\left(P_{e,v}P_{e,w}\right) + p^D\left(2P_{e,u}E_e\right) \\ & + p^T\left(\bar{P}^T_w P_{e,v} + \bar{P}^T_v P_{e,w}\right) + p^T\left(\bar{P}^T_u E_e + \bar{E}^T P_{e,u}\right), \end{aligned} \tag{7}$$

where:

$$\bar{P}^T_u = \sum_{h \in S} \frac{P_{h,u}}{|S|}, \tag{8}$$

Let $r$ be the gene family tree (GFT) root. Then:

$$P(G|S) = \sum_{s \in S} P_{r,s} \tag{9}$$

So far, we assumed that $G$ is rooted. If $G$ is unrooted, we can compute $P(G|S)$ by summing the likelihood score over all possible root locations (i.e., over all branches).

Let $\mathcal{G}$ be a set of GFTs and let $S$ be a species tree. The reconciliation likelihood is computed as:

$$L(S|\mathcal{G}) = \prod_{G \in \mathcal{G}} P(G|S) \tag{10}$$

Note that, over 95% of the overall species tree search runtime is spent in computing the reconciliation likelihood $P(G|S)$. The time complexity of the reconciliation function is $O(|S||G|)$, because it needs to compute the value $P_{e,u}$ for all branches $e \in S$ and $u \in G$. In the following sections, we describe how we reduce the computational cost of the reconciliation likelihood calculations during the tree search.

#### 1.2 The HGT-Loss approximation

We first observe that Eq. 7 is not an analytic formula but a system of equations, because the term for computing $P_{e,u}$ depends on itself and on $P_{h,u}$ for all nodes $h$ in the species tree. This is due to the duplication-loss term $p^D\left(2P_{e,u}E_e\right)$ and to the HGT-loss term $p^T\left(\bar{P}_u^T E_e + \bar{E}^T P_{e,u}\right)$. While the first term can be computed analytically, we need to deploy numerical optimization routines to evaluate the second term.

In our HGT-loss approximation approach, we simply discard this term from the initial likelihood formula. Thus, we do not to account for scenarios involving a gene $u$ being transferred from a species $e$ to a species $f$ and going extinct after $e$.

We ran the experiments for the current paper as well as the our previous experiments with GeneRax ([Morel *et al.*(2019)Morel, Kozlov, Stamatakis, and Szöllősi]) with and without this approximation. We did not observe any difference in species tree reconstruction accuracy for SpeciesRax and in GFT reconstruction accuracy for GeneRax. However, the reconciliation likelihood evaluation runs three times faster compared to the exact evaluation.

#### 1.3 Rooting the GFTs

The input GFTs are unrooted. Yet computing the reconciliation likelihood on a rooted GFT is substantially faster than for an unrooted GFT. First, this is because for unrooted GFTs, we need to iterate over all possible GFT root positions, and therefore need to compute every $P_{e,u}$ term three times (once for each possible orientation of the outgoing branches of the GFT node $u$ toward a potential root) instead of once for a single GFT root. Second, in the following subsection we introduce the *double-HGT approximation* to accelerate the computation of $P_{e,u}$ for GFT internal nodes $u$ that are far from the root (in terms of internode distance). Thus, iterating over all possible roots would substantially reduce the overall speedup that can be obtained via the double-HGT approximation.

For the above reasons, we only compute the likelihood for the maximum likelihood (ML) root position of each GFT. To infer this ML root, we perform a local GFT root search for each new species tree candidate as follows: Let us assume that we know the best root position of a GFT $G$ for a given species tree $S$. When evaluating the likelihood of a new species tree $S'$, we evaluate its value for the previously best GFT root position and for placing the root into the neighboring branches. Note that the additional cost for exploring the neighboring putative root positions is negligible for large GFTs, as the majority of the recursive intermediate computations are redundant for all five root positions. If one of the four putative neighboring root positions yields a better likelihood, we set it as the new best root. Then we repeat the above procedure on the four neighboring branches of this new root until no better root position is found. We omit the computations for the root position that we have already tested. Note that this local root search is not guaranteed to find the globally optimal ML root of the GFT.

Let $\mathcal{G}$ be a set of GFTs, let $S$ be the current species tree and let $S'$ be a new candidate species tree. Let $L(S, \mathcal{G})$ be the likelihood of $S$. By $L$ we denote the likelihood obtained for the globally optimal ML GFT root positions and by $\hat{L}$ we denote the likelihood of the best respective roots obtained via the local root search procedure described above. Obviously, $\hat{L}(S', \mathcal{G}) \leq L(S', \mathcal{G})$ because the local root search might not find the globally optimal ML roots on all GFTs. To approximately correct for this underestimation when comparing $S$ and $S'$, we test:

$$\hat{L}(S', \mathcal{G}) + \epsilon \geq L(S, \mathcal{G}) \tag{11}$$

where $\epsilon$ is twice the average underestimation for all previously accepted species trees $\Phi$:

$$\epsilon = 2\frac{\sum_{S \in \Phi} L(S, \mathcal{G}) - \hat{L}(S, \mathcal{G})}{|\Phi|} \tag{12}$$

When the test in Eq. 11 is positive, we exactly evaluate $L(S', \mathcal{G})$ via an exhaustive GFT root search for each gene family. We then accept $S'$ as the new current species tree if:

$$L(S', \mathcal{G}) > L(S, \mathcal{G}) \tag{13}$$

To initialize $\epsilon$, we skip the test in Eq. 11 for the 20 first candidate species trees. We determined this value (20) empirically via computational experiments on simulated and empirical datasets.

#### 1.4 The double-HGT approximation

Let $S$ be a rooted species tree and let $G$ be a rooted GFT. Computing the term $P(G|S)$ has time complexity $O(|S||G|)$ because it consists in evaluating $P_{e,u}$ for all species nodes $e$ of $S$ and all gene nodes $u$ of $G$. We observe that under the UndatedDTL model, the probability $\frac{P_T}{|S|}$ of an HGT event between two species is typically substantially smaller than the probabilities of other events ($P_S$, $P_L$, and $P_D$). Hence, unlikely HGT events highly penalize the reconciliation likelihood scores.

Further, we observe the following: an ancestral GFTs node $u$ is very unlikely to be observed in a species branch $e$ that is not the ancestor of at least one terminal species in which at least one GFT terminal node descending from $u$ is observed. This is because such a scenario would require at least two HGT events to be explained (one on each lineage of $u$) and would therefore penalize the reconciliation likelihood to a larger extent than an alternative scenario with one single HGT event prior to $u$.

Let $L(u)$ be the set of GFT terminal nodes that descend from $u$. Let $L_S(u)$ be the set of species that are mapped to the elements of $L(u)$. Let $X_u$ be the lowest common ancestor of $L_S(u)$ in $S$. In our approximation, we only compute $P_{e,u}$ if $e$ is either an ancestor or a descendant of $X_u$, and set $P_{e,u} := 0$ otherwise.

We do not attempt to formally estimate the speedup obtained via this approximation because it depends on the *G and S* tree topologies. We remark that the GFT nodes $u$ where $X_u$ is close to the leaves of the species tree will require few computations and that we can expect a large fraction of GFT nodes to be located close to the leaves as, in practice, most nodes in a binary rooted tree are closer to the leaves than to the root.

### 2 Tree search algorithm

The SpeciesRax search algorithm consists of four separate steps: ML species tree root inference, DTL intensities optimization, local subtree prune and regraft (SPR) species tree search and transfer-guided SPR species tree search. In this section, we first describe each of these four steps in details, and subsequently describe in which order we apply them.

#### 2.1 Maximum likelihood species tree root inference

To infer the ML root of a given species tree, SpeciesRax roots the species tree at several candidate positions, evaluates the reconciliation likelihood of each new putative root position, and keeps the best one. In the *exhaustive* root search step, SpeciesRax evaluates all possible putative root positions. In the *local* root search step, SpeciesRax only explores putative root positions around a given radius of the current root (typically, all branches that are less than three nodes away from the current root).

#### 2.2 DTL intensities optimization

SpeciesRax optimizes the DTL intensities via a gradient descent method. SpeciesRax provides two modes. In the *global* DTL intensities mode, all families share the same three (duplication, loss, and HGT) intensities. In the *per-family* DTL intensities mode, each gene family has its own set of DTL intensities to account for DTL rate heterogeneity. In our experiments we observed that the choice of the mode does not significantly affect runtime and that the per-family DTL intensities mode yields slightly improved species trees.

#### 2.3 Local SPR search

In the *local SPR search*, we explore all possible SPR moves for a given subtree rearrangement radius (i.e., the number of nodes away from the subtree pruning position at which we attempt to re-insert the subtree again). The default rearrangement radius is set to 1, because we observed on our experiments that higher values did not improve the reconstruction accuracy. We directly keep the trees generated by SPRs that improve the reconciliation likelihood. We stop this procedure when no better tree is found.

#### 2.4 Transfer-guided SPR search

In the *transfer-guided SPR search*, we assume that the most promising SPR moves with respect to improving the reconciliation likelihood are those SPR moves that reduce the number of horizontal gene transfer (HGT) events that are necessary to reconcile the GFTs with the species tree (a similar strategy has been previously applied in [Szöllősi *et al.*(2012)Szöllősi, Boussau, Abby, Tannier, and Daubin]). To this end, we infer the ML reconciliation between the GFT and the current species tree, and count the number of HGTs between each pair of nodes in the species tree. We then try the SPR moves between the pair of species (we regraft the node of the species tree corresponding to the receiving species lineage next to the node of the species tree corresponding to the source species lineage) that yields the highest numbers of HGTs, and apply those SPR moves that improve the reconciliation likelihood if any. We stop these attempts after $n$ unsuccessful consecutive trials, where $n$ is the number of species. After $k = max(15, n/4)$ non-consecutive successful trials, we re-infer the HGTs on the current species tree. We empirically determined the best values of $n$ and $k$.

#### 2.5 Species tree search overview

We now explain how we combined the previously described moves to infer a rooted ML species tree.

If the starting species tree was provided by the user or generated with MiniNJ, we optimize the DTL rates and execute a local root search. If the starting species tree was randomly generated, we start from an predefined set of rates ($\delta = \tau = \lambda = 0.2$). Then we apply the transfer-guided and local SPR searches in an alternating manner. After each (transfer-guided or local) SPR search, we conduct a local species tree root search and optimize the DTL rates. When no better species tree can be found, we run a final local species tree root search with a higher radius (5 by default instead of 3 for the previous ones) and stop the search. The aforementioned *exhaustive* species tree root search is optional and is not executed by default.

### 3 Species tree branch length estimation

SpeciesRax infers the branch lengths of the rooted species tree in units of expected number of substitutions per site. We assume that the GFTs are reconciled with the species tree and have their branch length in units of substitution per site. Our method independently estimates the length of each species branch, averaging over the branch lengths between relevant speciation events in the reconciled GFTs.

Remember that both, the species tree, and the reconciled GFTs are rooted. For any rooted tree $T$, let $N(T)$ be the set of nodes in $T$. For any $x \in N(T)$, let $t(x)$ be the branch length of $x$. Let $S$ be a species tree, let $\mathcal{G}$ be the set of GFTs, and let $G \in \mathcal{G}$. Let $(s, \sigma)$ be the reconciliation function that maps each gene node $x \in G$ to a species node $s(x) \in N(S)$ and to an event label

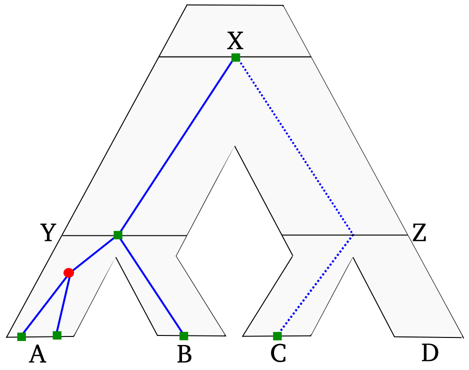


Figure 1: Illustration of relevant paths for the species tree branch length estimation. A GFT (in blue) is represented within a species tree (tree with grey background). Relevant paths are represented by solid lines, and non-relevant paths by dashed lines. The gene tree has one $Y$-relevant path, one $B$-relevant path and two $A$-relevant paths (due to the duplication event along $A$). The gene branch that leads from species $X$ to species $C$ is not part of a relevant path because $C$ is not a direct child of $X$: this branch goes through an unobservable speciation (the gene goes extinct in the branch of species $D$) and thus the time of the speciation at $Z$ is unknown.

$\sigma(x) \in \{E_S, E_D, E_T\}$ (speciation, duplication, and HGT). We define a *path* $p$ in $G$ as a sequence of nodes in $N(G)$ such that each element in $p$ is the child of its predecessor. We define the length $t(p)$ of a path $p$ as the sum of the lengths of its elements.

We now introduce the concept of *$f$-relevant paths* to characterize gene paths that contain relevant information for estimating the branch length above a species node $f \in N(S)$. For a given $f \in N(S)$ and its parent node $e$, an *$f$-relevant path* $p = (x_1, x_2, ..., x_{|p|})$ in $G$ is a path such that $\sigma(x_1) = \sigma(x_{|p|}) = E_S$, $s(x_1) = e$ and $s(x_{|p|} = f)$ (See example in Fig. 1). By $\mathcal{P}_f(g)$ we denote the set of all $f$-relevant paths in $G$.

For each species node $f$, we compute its length as the weighted average of all $f$-relevant paths:

$$\hat{t}(f) = \frac{\sum_{G \in \mathcal{G}} \sum_{p \in \mathcal{P}_f(G)} w(G) t(p)}{\sum_{G \in \mathcal{G}} \sum_{p \in \mathcal{P}_f(G)} w(G)}$$

where $w(G)$ is a weight function associated to each GFT. If the multiple sequence alignment (MSA)s are available, we set $w(G) = l_G r_G$, where $l_G$ is the length of the MSA associated with $G$, and $r_G$ is the proportion of characters thar are neither undetermined nor gaps. If the MSAs are not available, we set $w(G) = 1$.

### 4 Simulation parameters

#### 4.1 SimPhy parameters

We summarize all SimPhy simulation parameters in Table 1.

#### 4.2 Justification for the DTL rates used in the simulations

In this section, we assess if the DTL rates $\lambda$ and $\tau$ (defined in Table 1) used in our simulations yield realistic numbers of DTL events (gene duplication, gene loss and GFT). To this end, we inferred the number of DTL events on a subset of the empirical and simulated datasets used in our benchmark. We ran GeneRax with its default parameters to correct the GFTs from the species trees and to compute the total number of speciations, duplications, losses, and HGTs. We applied the procedure to the empirical datasets primates13, cyanobacteria36, vertebrates21, fungi59, and plants23. We did not apply it to fungi16 and plants83 because their gene sequences were not available. We did also not apply it to vertebrates188 because it was computationally too expensive. On the simulated datasets, we applied the procedure to 10 replicates generated with default parameters under DLSIM and D**T**LSIM, respectively.

We summarize the results in Table 2 and Table 3, respectively. We also calculated the ratio between inferred duplication and speciation events. We observe that this ratio as inferred for DLSIM lies within the range of ratios inferred on the empirical datasets (0.27 for DLSIM, 0.10 for primates, 0.26 for vertebrates, and 0.37 for plants). The same applies to the ratio between inferred HGT and speciation events in the simulations with DTL events (0.12 for D**T**LSIM, 0.13 for cyanobacteria and 0.10 for fungi). This suggests that the duplication, HGT, and speciation rates we used for our simulations are realistic and match those of empirical datasets.

However, we seem to always underestimate the frequency of loss events, although this quantity is harder to infer: one can not determine if inferred loss events correspond to actual gene loss events or are due to missing data or inaccurate gene family clustering. Unfortunately, SimPhy does not allow to set a loss rate that is higher than the duplication rate.

### 5 Species tree RF-distance Table

In Table 4, for each empirical dataset, we provide the RF distances between all pairs of inferred species trees.

### References

[Morel *et al.*(2019)Morel, Kozlov, Stamatakis, and Szöllősi] Morel, B., Kozlov, A. M., Stamatakis, A., and Szöllősi, G. J. 2019. Generax: A tool for species

| Parameter name | Parameter value |
|---|---|
| Standard parameters | |
| Replicates number | 100 |
| Speciation rate | $5 \times 10^{-9}$ |
| Extinction rate | $4.9 \times 10^{-9}$ |
| Number of gene families | 100 |
| Number of species | 25 |
| Dup and loss rates | $\delta \times Log\text{-}\mathcal{N}(0,1)$, $\delta = 4.9 \times 10^{-10}$ |
| HGT rate | $\tau \times Log\text{-}\mathcal{N}(0,1)$, $\tau = 4.9 \times 10^{-10}$ |
| Population size | 10 |
| Species tree height | $Log\text{-}\mathcal{N}(21.25, 0.2)$ |
| Global substitution rate | $Log\text{-}\mathcal{N}(-21.9, 0.1)$ |
| Lineage specific rate gamma shape | $Log\text{-}\mathcal{N}(1.5, 1)$ |
| Family specific rate gamma shape | $Log\text{-}\mathcal{N}(1.551533, 0.6931472)$ |
| Gene tree branch specific rate gamma shape | $Log\text{-}\mathcal{N}(1.5, 1)$ |
| Sequence length | $\nu \times Log\text{-}\mathcal{N}(0, 0.25)$, $\nu = 100(e^{-\frac{0.25^2}{2}})$ |
| Sequence base frequencies | Dirichlet(A=36,C=26,G=28,T=32) |
| Sequence transition rates | Dirichlet(TC=16,TA=3,TG=5,CA=5,CG=6,AG=15) |
| Seed | $[3000, 3100[$ |
| Varying parameters | |
| Dup and loss rates multiplier | 0.5,1.0,2.0,3.0 |
| HGT rates multiplier | 0, 0.5, 1, 2.0, 4.0 |
| Population size | 10, $10^7$, $10^8$, $10^9$ |
| Number of species | 15, 25, 35, 50, 75 |
| Number of gene families | 50, 100, 200, 500, 1000 |
| Average number of sites | 50, 100, 200, 300 |

Table 1: SimPhy parameters to simulate the SIMDL and SIMDTL datasets. In the varying parameters section, the rate multipliers are used to scale the constants $\lambda$ for the dup-loss rates and $\tau$ for the HGT rates. For sequence length, $\nu$ is set to obtain 100 sites on average.

| Dataset | S | D | L | D/S |
|---|---|---|---|---|
| DLSIM | 0.72 | 0.19 | 0.09 | 0.27 |
| Primates13 | 0.75 | 0.08 | 0.18 | 0.10 |
| Vertebrates21 | 0.59 | 0.15 | 0.26 | 0.26 |
| Plant23 | 0.55 | 0.20 | 0.25 | 0.37 |

Table 2: Frequencies of speciation (S), duplication (D) and loss (L) events inferred with GeneRax on various datasets. DLSIM corresponds to the average over 10 replicates of the datasets generated under the default parameter set of the DLSIM set of simulations.

| Dataset | S | D | L | T | T/S |
|---|---|---|---|---|---|
| D**T**LSIM | 0.69 | 0.15 | 0.08 | 0.09 | 0.12 |
| Cyanobacteria36 | 0.80 | 0.02 | 0.08 | 0.11 | 0.13 |
| Fungi59 | 0.74 | 0.04 | 0.14 | 0.08 | 0.10 |

Table 3: Frequencies of speciation (S), duplication (D) loss (L), and HGT (T) events inferred with GeneRax on various datasets. D**T**LSIM frequencies corresponds to the average over 10 replicates on the datasets generated under the default parameter set of the D**T**LSIM simulations.

| | S | M | A | F | D |
|---|---|---|---|---|---|
| S | 0 | 0 | 0 | 0 | 0 |
| M | 0 | 0 | 0 | 0 | 0 |
| A | 0 | 0 | 0 | 0 | 0 |
| F | 0 | 0 | 0 | 0 | 0 |
| D | 0 | 0 | 0 | 0 | 0 |

(a) primates13

| | S | M | A | F | D |
|---|---|---|---|---|---|
| S | 0 | 2 | 0 | 0 | 0 |
| M | 2 | 0 | 2 | 2 | 2 |
| A | 0 | 2 | 0 | 0 | 0 |
| F | 0 | 2 | 0 | 0 | 0 |
| D | 0 | 2 | 0 | 0 | 0 |

(b) cyanobacteria36

| | S | M | A | F | D |
|---|---|---|---|---|---|
| S | 0 | 0 | 0 | 2 | 4 |
| M | 0 | 0 | 0 | 2 | 4 |
| A | 0 | 0 | 0 | 2 | 4 |
| F | 2 | 2 | 2 | 0 | 2 |
| D | 4 | 4 | 4 | 2 | 0 |

(c) vertebrates21

| | S | M | A | F | D |
|---|---|---|---|---|---|
| S | 0 | 2 | 0 | 0 | 2 |
| M | 2 | 0 | 2 | 2 | 4 |
| A | 0 | 2 | 0 | 0 | 2 |
| F | 0 | 2 | 0 | 0 | 2 |
| D | 2 | 4 | 2 | 2 | 0 |

(d) fungi59

| | S | M | A | F | D |
|---|---|---|---|---|---|
| S | 0 | 2 | 0 | 0 | 2 |
| M | 2 | 0 | 2 | 2 | 4 |
| A | 0 | 2 | 0 | 0 | 2 |
| F | 0 | 2 | 0 | 0 | 2 |
| D | 2 | 4 | 2 | 2 | 0 |

(e) fungi60

| | S | M | A | F | D |
|---|---|---|---|---|---|
| S | 0 | 14 | 6 | 6 | 2 |
| M | 14 | 0 | 12 | 8 | 16 |
| A | 6 | 12 | 0 | 8 | 4 |
| F | 6 | 8 | 8 | 0 | 8 |
| D | 2 | 16 | 4 | 8 | 0 |

(f) plants23

| | S | M | A | F | D |
|---|---|---|---|---|---|
| S | 0 | 18 | 14 | 18 | 60 |
| M | 18 | 0 | 10 | 10 | 66 |
| A | 14 | 10 | 0 | 6 | 60 |
| F | 18 | 10 | 6 | 0 | 62 |
| D | 60 | 66 | 60 | 62 | 0 |

(g) plants83

| | S | M | A | F | D |
|---|---|---|---|---|---|
| S | 0 | 12 | 14 | 8 | 54 |
| M | 12 | 0 | 10 | 12 | 52 |
| A | 14 | 10 | 0 | 6 | 52 |
| F | 8 | 12 | 6 | 0 | 54 |
| D | 54 | 52 | 52 | 54 | 0 |

(h) vertebrates188

Table 4: RF distances between empirical species trees inferred with the tested tools: SpeciesRax (S), MiniNJ (M), ASTRAL-Pro (A), FastMulRFS (F) and DupTree (D).

tree-aware maximum likelihood based gene tree inference under gene duplication, transfer, and loss. *bioRxiv*.

[Szöllősi *et al.*(2012)Szöllősi, Boussau, Abby, Tannier, and Daubin] Szöllősi, G. J., Boussau, B., Abby, S. S., Tannier, E., and Daubin, V. 2012. Phylogenetic modeling of lateral gene transfer reconstructs the pattern and relative timing of speciations. *Proceedings of the National Academy of Sciences*, 109(43): 17513–17518.
